# Parameterized syncmer schemes improve long-read mapping

**DOI:** 10.1101/2022.01.10.475696

**Authors:** Abhinav Dutta, David Pellow, Ron Shamir

## Abstract

**Motivation:** Sequencing long reads presents novel challenges to mapping. One such challenge is low sequence similarity between the reads and the reference, due to high sequencing error and mutation rates. This occurs, e.g., in a cancer tumor, or due to differences between strains of viruses or bacteria. A key idea in mapping algorithms is to sketch sequences with their minimizers. Recently, syncmers were introduced as an alternative sketching method that is more robust to mutations and sequencing errors.

**Results:** We introduce parameterized syncmer schemes, a generalization of syncmers, and provide a theoretical analysis for multi-parameter schemes. By combining these schemes with downsampling or minimizers we can achieve any desired compression and window guarantee. We implemented the use of parameterized syncmer schemes in the popular minimap2 and Winnowmap2 mappers. In tests on simulated and real long read data from a variety of genomes, the syncmer-based algorithms, with scheme parameters selected on the basis of the theoretical analysis, reduced unmapped reads by 20-60% at high compression while usually using less memory. The advantage was more pronounced at low sequence identity. At sequence identity of 75% and medium compression, syncmer-minimap had only 37% as many unmapped reads, and 8% fewer of the reads that did map were incorrectly mapped. Even at lower compression and error rates, parameterized syncmer based mapping mapped more reads than the original minimizer-based mappers as well as mappers using the original syncmer schemes. We conclude that using parameterized syncmer schemes can improve mapping of long reads in a wide range of settings.

**Availability:** https://github.com/Shamir-Lab/syncmer_mapping

**Supplementary information:** Supplementary data are available at https://github.com/Shamir-Lab/syncmer_mapping.

**Author summary:** Popular long read mappers use minimizers, the minimal hashed *k*-mers from overlapping windows, as alignment seeds. Recent work showed that syncmers, which select a fixed set of *k*-mers as seeds, are more likely to be conserved under errors or mutations than minimizers, making them potentially useful for mapping error-prone long reads. We introduce a framework for creating syncmers, that we call parameterized syncmer schemes, which generalize those introduced so far, and provide a theoretical analysis of their properties. We implemented parameterized syncmer schemes in the minimap2 and Winnowmap2 long read mappers. Using parameters selected on the basis of our theoretical analysis we demonstrate improved mapping performance, with fewer unmapped and incorrectly mapped reads on a variety of simulated and real datasets. The improvements are consistent across a broad range of compression rates and sequence identities, with the most significant improvements for lower sequence identity (high error or mutation rates) and high compression.

## Introduction

As the volume of third-generation, long read sequencing data increases, new computational methods are needed to efficiently analyze massive datasets of long reads. One of the most basic steps in analysis of sequencing data is mapping reads to a known reference sequence or to a database of many sequences. Several long read mappers have been proposed [1,2], with minimap2 [3] being the most popular. minimap2 is a multi-purpose sequence mapper that uses sequence minimizers as alignment anchors. Minimizers, the minimum valued *k*-mers in windows of *w* overlapping *k*-mers of a sequence, are used to sketch sequences. They have greatly improved the computational efficiency of many different sequence analysis algorithms (e.g. [4], [5], [6]). A key criterion in evaluating minimizer schemes is their *compression rate*, the number of *k*-mers in the sequence divided by the number of *k*-mers selected. Achieving higher compression rate is desirable, as fewer anchors are used.

Recent work has shown that minimizers are less effective under high error or mutation rates [7]. Motivated by this observation, Edgar [7] recently introduced a novel family of *k*-mer selection schemes called *syncmers*. Syncmers are a set of *k*-mers defined by the position of their minimum *s*-long substring (*s*-minimizer). Syncmers constitute a predetermined subset of all possible *k*-mers and, unlike minimizers, they are defined by the sequence of the *k*-mer only and do not depend on the rest of the window in which they appear. Syncmers are therefore more likely to be conserved under mutations than minimizers. This difference is crucial in long reads, which have much higher error rate than short reads [8]. Another key difference between syncmer and minimizer schemes is that the latter guarantee selection of a *k*-mer in every window of *w* consecutive *k*-mers (this is called a *window guarantee*), while syncmers do not. For longer reads with a higher error rate, *conservation* of the selected *k*-mers becomes more important than the window guarantee, especially when there are also mutations. For example, it was shown that with 90% identity between aligned sequences, only about 30% of the positions on the sequence will overlap a conserved minimizer in minimap2 [7].

Edgar defined several syncmer variants, including the families of *open syncmers*, whose *s*-minimizer appears at one specific position, and *closed syncmers*, whose *s*-minimizer appears at either the first or the last position in the *k*-mer [7]. He computed the properties of a range of syncmer schemes and used them to choose a scheme with a desired compression rate. Shaw and Yu [9] recently formalized the notions of the conservation of selected positions and their clustering along a sequence, and provided a broader theoretical analysis.

In this work we generalize Edgar’s syncmer schemes to multiple arbitrary *s*-minimizer positions. We call these *parameterized syncmer schemes* (PSS; we use this acronym for both singular and plural). The parameters are the possible indices of the *s*-minimizer in a selected *k*-mer, and an *n*-parameter scheme uses *n* such indices. An example is a 3-parameter scheme that selects any 15-mer with the minimum 5-mer appearing at position 1, 5, or 9. We analyze the properties of such parameterized syncmer schemes and determine which schemes perform well for a given compression rate through theoretical analysis and empirical testing. PSS have the advantage of allowing for a larger range of compression rates than syncmers, by varying the number of parameters used, *k*-mer length, and *s*-minimizer length. Additionally, while closed syncmers are a subset of 2-parameter PSS, our analysis demonstrates that they are not the optimal 2-parameter scheme under realistic mutation rates and enables us to instead select the best 2-parameter scheme.

Two important related features of a scheme are robustness to sequence changes and the distances between selected positions. The conservation of a scheme is the fraction of positions in a sequence covered by selected *k*-mers that are unchanged after the sequence is mutated. The spread of a scheme is a vector of probabilities, where *P*(*α*) is the probability of selecting at least one position in a window of length *α*. Recently, Shaw and Yu [9] obtained expressions for the conservation of open and closed syncmers as a function of spread and implemented these syncmers in minimap2. Here we extend the theoretical analysis by presenting a general recursive expression for the spread of any PSS, including downsampling. These expressions allow for the calculation of the conservation of any PSS.

We introduced PSS into two leading long read mappers: the latest release of minimap2 [10] and Winnowmap2 [11], where PSS parameters were selected based on our theoretical analysis, and measured the performance compared to the original algorithms on both simulated and real long read data. The syncmers increased the number of mapped reads across a large range of compression rates, resulting in 20-60% fewer unmapped reads at high compression. Even at lower compression, the syncmer mappers had 2-15% fewer unmapped reads. The syncmer versions used less memory but had longer mapping times than the original mappers for the same compression. The most marked improvements were observed when identity of the mapped reads and reference sequences were low. With identity of 65% and 75% and medium compression, syncmer mappers had 50-60% fewer unmapped reads and still had 8-13% fewer incorrectly mapped reads. When using the best 2-parameter PSS according to our theoretical analysis in minimap2 in comparison to minimap2 using Edgar’s 2-parameter closed syncmers, the former showed a consistent improvement of up to 7% fewer unmapped reads.

Our contributions in this work are thus three-fold: (1) We introduce PSS, generalizing existing syncmer schemes. (2) We provide a theoretical analysis of PSS properties. The analysis enables us to choose the optimal scheme for particular mutation and compression rates. (3) We provide implementations of minimap2 and Winnowmap2 that use PSS and demonstrate their improved mapping performance compared to the original minimizer versions and to using open syncmers. Unlike previous work, our implementations also enable downsampling, so that any desired compression rate can be achieved, and on the other hand they have the option to provide a window guarantee.

The paper is structured as follows: we first provide background, definitions, and terminology; the next section provides theoretical analysis of PSS and describes the practical implementation of PSS and their integration into minimap2 and Winnowmap2; the following section presents experimental results of the original and PSS-modified mappers; the final section discusses the results and future work.

## Definitions and Background

### Basic definitions and notations

For a string *S* over the alphabet Σ, a *k*-mer is a *k*-long contiguous substring of *S*. The *k*-mer starting at position *i* is denoted *S*[*i, i* + *k* – 1] (string indices start from 1 throughout). We work with the nucleotide alphabet: Σ = {*A, C, G, T*}.

#### *k*-mer order

Given a one-to-one hash function on *k*-mers *o*: 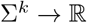, we say that *k*-mer *x*_1_ is less than *x*_2_ if *o*(*x*_1_) < *o*(*x*_2_). Examples include lexicographic encoding or random hash. We will write instead *x*_1_ < *x*_2_ when o is clear from the context. In this work we use a random order unless otherwise noted.

#### Canonical *k*-mers

Denote the reverse complement of *x* by 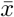. For a given order, the *canonical form* of a *k*-mer *x*, denoted by *Canonical*(*x*), is the smaller of *x* and 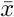. For example, under the lexicographic order, *Canonical*(*CGGT*) = *ACCG*.

### Selection schemes

#### Selection scheme

A *selection scheme* is a function from a string to the indices of positions in it 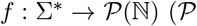 represents the power set). The scheme implicitly selects the *k*-mers starting at these positions. For a string *S* ∈ Σ*, *f_k_*(*S*) = {*i*_1_, *i*_2_,…, *i_n_*} is the set of start indices of the *k*-mers selected by the scheme.

#### Minimizers

A *minimizer scheme* is a selection scheme that chooses the position of the minimum value *k*-mer in every window of *w* consecutive *k*-mers in *S*:

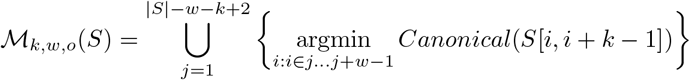

where the minimum is according to *k*-mer ordering *o*. By convention, ties are broken by choosing the leftmost position. An example of a minimizer selection scheme is shown in Fig 1A. By definition, minimizers select a *k*-mer in every window of *w k*-mers. This property is called a *window guarantee*.

**Fig 1.**
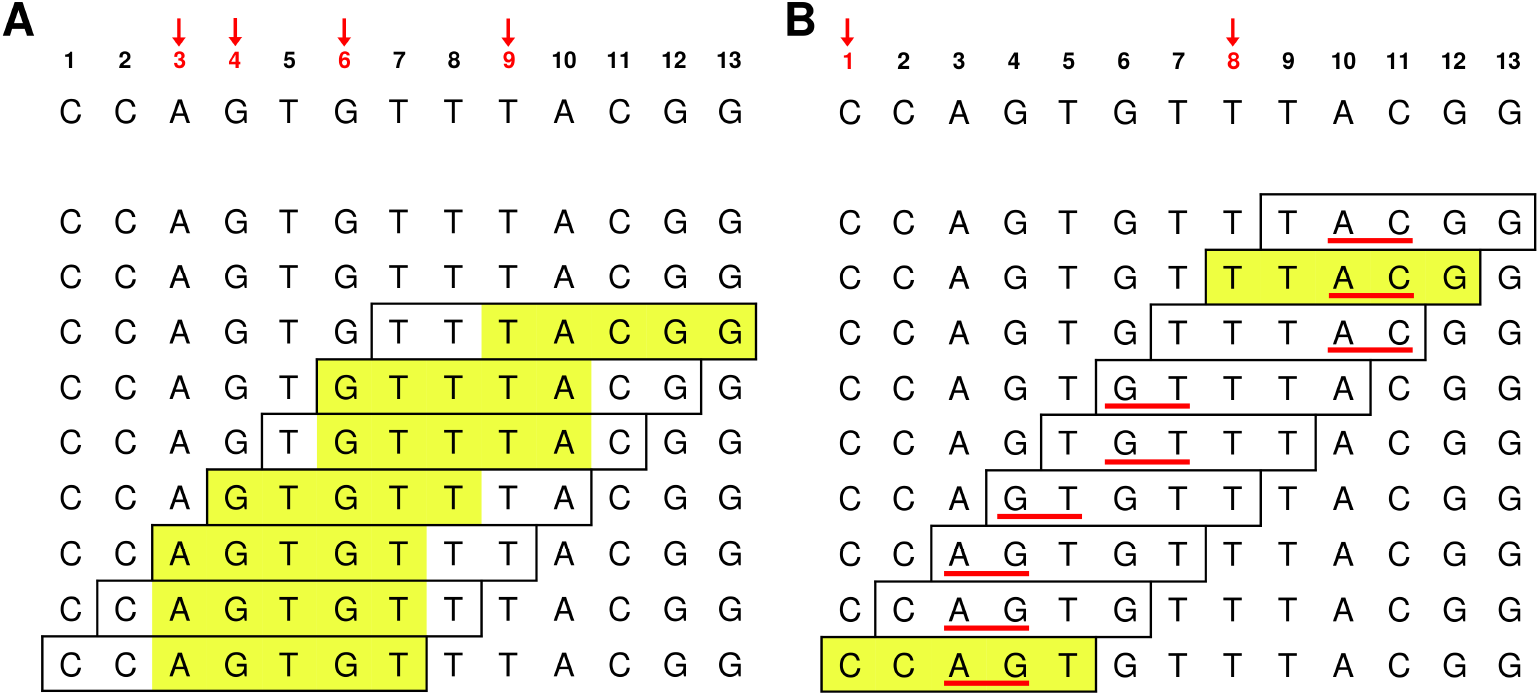
Minimizer and syncmer schemes. In both examples the lexicographic order is used and the underlying sequence is shown at the top. By convention the leftmost position is selected in the case of a tie. **(A)** Minimizers. Here *w* = 3 and *k* = 5, so the minimizer is the least 5-mer in every window of length 7. The minimizer of each window is highlighted in yellow; **(B)** Syncmers. Here we show the 1-parameter syncmer with *k* = 5, *s* = 2 and *t* = 3, 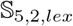(3). It selects 5-mers if their 2-minimizer appears at position 3. The 2-minimizer in each 5-mer is underlined in red, and selected *k*-mers are highlighted in yellow. The start positions of the *k*-mers in the underlying sequence that are selected by each scheme appear in red and are marked with red arrows at the top. Sequence positions 6-7 constitute a gap in the syncmer selection as they are not covered by any selected *k*-mer.

#### Syncmers

A *syncmer* [7] is a selection scheme that selects a *k*-mer if its minimum *s*-mer is in a particular position or positions. A *closed syncmer* selects *k*-mers whose smallest *s*-mer is at the start or end of the *k*-mer, and an *open syncmer* select *k*-mers whose smallest *s*-mer is at the start only. Note that, unlike minimizers, a syncmer does not have a window guarantee since it will only identify a fixed subset of all *k*-mers

#### Parameterized syncmers

We are now ready to introduce the key new concept of this study. A *parameterized syncmer scheme* (PSS) with parameters *s, k, o* and *x*_1_,…, *x_n_* where 0 < *x*_*n*-1_ <… <*x*_*n*-1_ < *x_n_* ≤ *k* – *s* + 1 selects a *k*-mer if the minimum *s*-mer of that *k*-mer appears at one of the positions *x_i_* in the *k*-mer. Formally:

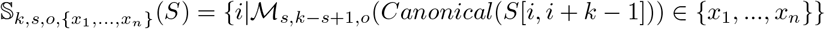

As *o* is fixed we will drop it from the notation where possible. An example of a PSS is shown in Fig 1B. For convenience, we will denote the syncmer scheme with parameters *x*_1_,…*x_n_* as 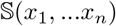 and the family of all *n*-parameter syncmers as 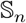. Under these definitions, the open and closed syncmer schemes are 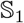 and 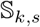(1, *k* – *s* + 1), respectively. From here on, we will refer to PSS simply as syncmers.

#### Downsampled and windowed syncmers

In some situations we wish to cull the selected *k*-mers or fill-in sequence segments where none was selected. Given a uniformly random hash function *h*: Σ^*k*^ → [0, *H*], for a given string *S*, *downsampling* selects syncmers only from the set of |Σ|^*k*^/*δ k*-mers with the lowest hash values. We call *δ* the *downsampling rate*. Windowed syncmers fill in gaps using a minimizer scheme, thus providing a window guarantee. See Supplement section S1 for formal definitions of these sets.

### Properties and evaluation criteria of schemes

We define some metrics for evaluating the performance of selection schemes.

#### Density and compression

The *density* of a scheme [12] is the expected fraction of positions selected by the scheme in an infinitely long random sequence: 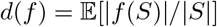 as |*S*| → ∞. The *compression rate* [7] is defined as *c*(*f*) = 1/*d*(*f*), i.e. the factor by which the sequence *S* is “compressed” by representing it using only the set of selected *k*-mers.

#### Conservation

Conservation [9] is the expected fraction of positions covered by a selected *k*-mer in sequence *S* that will also be covered by the same selected *k*-mer in the mutated sequence *S*’ where *S*’ is generated by iid base mutations with rate *θ*. Define the set of positions covered by the same selected *k*-mer in both sequences

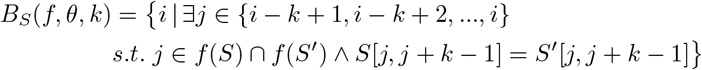

Then the *conservation* of the scheme is defined as 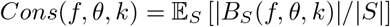.

#### Spread and distance distribution

One key feature of a scheme is the distance between selected positions and the frequency with which selected positions appear close together or far apart. Shaw and Yu [9] studied the probability distribution of selecting *at least one* position in a window of length *α*. We refer to the vector *P*(*f, α*) of these probabilities as the *spread*.

We define the *distance distribution* of *consecutive* selected positions: *Pr*(*f, n*) is the probability that position *i* + *n* is the next selected position given that position *i* is selected.

#### *pN* metric

The *pN* metric (*N* ∈ [0,100]) is the *Nth* percentile of the distance distribution, i.e., it is the length *l* for which *N*% of the distances between consecutive selected positions are of length ≤ *l*.

#### *ℓ* and *ℓ*_2_ metrics

A *gap* is a nonempty stretch of sequence between two consecutive selected *k*-mers. Gaps are uncovered by the scheme. Let the lengths of the gaps generated by a scheme on the sequence *S* be *l*_1_, *l*_2_,…. We define 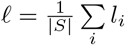 and 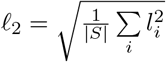.

While these metrics are defined in expectation for given sequence and mutation models and the selection scheme, we also use the analogously defined empirical values measured on a specific sequence. The metrics may also be considered on the positions selected by a scheme in a reference, or only on the selected positions that are *conserved* after mutation or sequencing error. We refer to the latter using the subscript *mut*, for example, *ℓ*_2,*mut*_ is defined analogously to *ℓ*_2_ except the gaps are between consecutive selected *k*-mers that are conserved after mutation.

### Choosing an appropriate metric to compare schemes

While Edgar shows convincingly that conservation is a more appropriate metric for selection schemes than density, we argue that *ℓ*_2,*mut*_ contains additional important information for the purpose of mapping. Specifically, observe that, for given mutation rate *θ*, *k*, and selection scheme *f*, we have 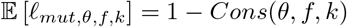. While *ℓ* (and conservation) counts the number of bases that are not covered by conserved selected *k*-mers, it treats all gap lengths equally. In contrast, *ℓ*_2,*mut*_ penalizes a few large gaps more than many smaller gaps with the same total length. See the example in Fig 2. When the selected *k*-mers are used as seeds for mapping, it is important to avoid large gaps, in order to enable read mapping across gaps. Thus *ℓ*_2_ provides additional information on how the selection scheme will affect mapping performance, and we use it throughout to select the scheme for our syncmer based mappers.

**Fig 2.**
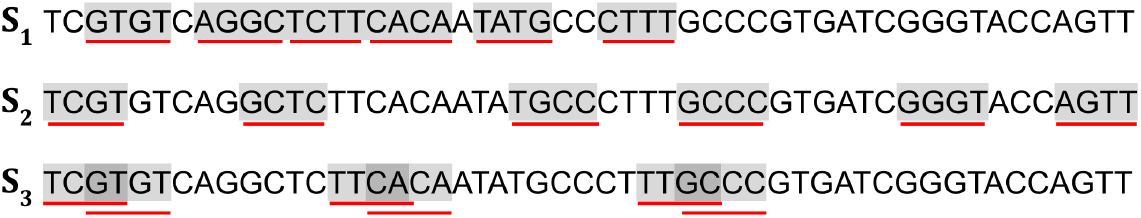
*ℓ* vs. *ℓ*_2_ metric. The selected positions of three different selection schemes *S*_1_, *S*_2_ and *S*_3_ on the same sequence. Selected *k*-mers are highlighted and underlined. All schemes have the same number of selected *k*-mers, but the metrics are different. **S**_1_: *ℓ* = 0.529, *ℓ*_2_ = 2.974. **S**_2_: *ℓ* = 0.529, *ℓ*_2_ = 1.81. **S**_3_: *ℓ* = 0.647, *ℓ*_2_ = 2.808. While S_1_ and S_2_ have the same *ℓ* value, the *k*-mers selected by *S*_2_ are more evenly spread and thus it has much lower *ℓ*_2_. Some of the *k*-mers selected by S_3_ overlap, resulting in a higher *ℓ* value than the other schemes. However, because the gaps between covered bases are more evenly spread, the *ℓ*_2_ value is lower than that of S_1_. Intuitively, it will be easier to map reads using seeds selected by S_3_ than S_1_ despite the higher *ℓ* value, suggesting that *ℓ*_2_ is a more appropriate metric.

### Analysis of syncmer schemes – prior work

Edgar recently defined syncmers as an alternative to minimizers and other selection schemes with the goal of improving conservation rather than density, arguing that conservation is often dictated by the application and system constraints [7]. He introduced open and closed syncmers and their rotated variants. He provided analyses of syncmer densities, window guarantees, and distributions were provided for open, closed, and downsampled syncmers.

Shaw and Yu greatly extended the framework for theoretical analysis of syncmers [9]. They defined the spread and conservation of a scheme. The two are connected through the number of unmutated *k*-mers overlapping a given position, *α*(*θ, k*), for a given mutation rate, *θ*. Letting *P*(*f*) = [*P*(*f*, 1), *P*(*f*, 2),… *P*(*f, k*)] be the spread, and *P*(*α*(*θ, k*)) = [*P*(*α*(*θ, k*) = 1),*P*(*α*(*θ, k*) = 2),…, *P*(*α*(*θ, k*) = *k*)], then *Cosns* (*f, θ, k*) = *P*(*f*) · *P*(*α*(*θ, k*)). Note that there is a closed form expression for calculating *P* (*α* (*θ, k*) = *α*)), and that *P*(*f*, 1) = *d*(*f*). Their theoretical framework allowed Shaw and Yu to obtain expressions for the spread (and therefore conservation) of open and closed syncmers and other selection schemes.

## Methods

In this section we first outline the main results of our theoretical analysis of PSS. These results provide guidance for choosing PSS parameters in practice. We then describe how we modify mappers to utilize them. Due to space constraints the full derivations and analysis are deferred to Supplementary section Analysis of parameterized syncmer schemes. Raw data for results presented in this and subsequent sections are available in Supplementary Data File 1 from https://github.com/Shamir-Lab/syncmer_mapping.

### Recursive expressions for conservation of PSS

Shaw and Yu [9] obtained expressions for open and closed syncmer conservation as a function of spread. Here we present a general recursive expression for the spread of any PSS, including with downsampling. These expressions allow for the calculation of the conservation of any PSS. The full derivation of the general expression is presented in Supplementary section S2.1 while here we present only the final expression itself.

Consider a window of *α* consecutive *k*-mers. We assume random sequence (i.e., made of iid bases) throughout. Let *s_β_* be the *s*-minimizer in the window, at position *β*. Then if *t* is a parameter of the syncmer scheme, *s_β_* generates a syncmer if it is not in the first *t* – 1 or last *k* – *t* positions in the window. If *β* is not in a position where it generates a syncmer, we recursively check to the left or right of *β* to see if a syncmer is generated by the *s*-minimizer of that region. See Fig 3 for an example.

**Fig 3.**
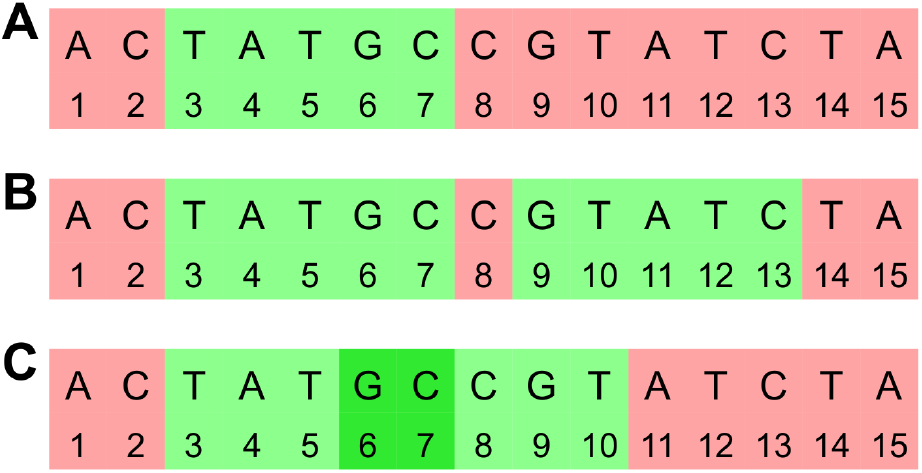
Illustration of *s*-minimizers generating syncmers. A window of *α* = 5 consecutive 11-mers. **A**: When *s* = 5 and *t* = 3, then the *s*-minimizer of the entire window generates a syncmer when its starting index is in the green region. If the *s*-minimizer is in one of the red regions then a syncmer may be generated by the *s*-minimizer of the remaining part of the window. For a two parameter scheme the *s*-minimizer creates two syncmer generating regions that may be disjoint (**B**) if *s* > *t*_2_ – *t*_1_ or overlapping (**C**) if *s* < *t*_2_ – *t*_1_. In this example, *t*_1_ = 3 and *t*_2_ = 9 in **B** and *t*_2_ = 6 in **C**.

For a PSS *f* with *k*-mer length *k, s*-minimizer length *s*, and downsampling rate *δ*, let *P*(*α*) be the probability of selecting at least one syncmer in a window of *α* adjacent *k*-mers. We assume a uniformly random hash over the *s*-mers, and condition on the position of the *s*-minimizer in the *α*-window, *β*. For each *β* we sum over two cases: 1) *β* generates at least one syncmer that is not lost due to downsampling, 2) *β* does not generate a syncmer, or all are lost due to downsampling, in which case a syncmer may be generated by the part of the window to the left or right, resulting in a recursive expression. Let *P_R_* = *P*(*α* – *β*) and *P_L_* = *P*(*β* – *k* + *s* – 1). Then we have:

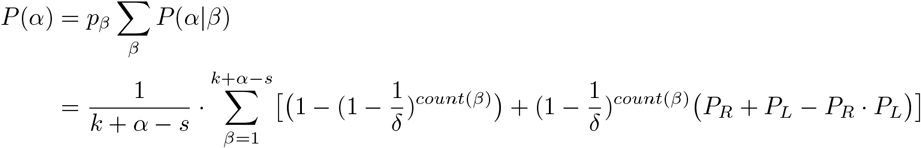

The probability of any of the *k* + *α* – *s* starting positions being the *s*-minimizer is denoted as *p_β_* and assumed to be uniform. *count*(*β*) represents the number of syncmers generated by the *s*-minimizer *s_β_*. For example, *count*(*β*) = 0 in the red region of Fig 3 and *count*(*β*) = 2 in the overlapped region when *β* = 6 or 7 in Fig 3C. Note we define *P*(*α*) = 0 when *α* ≤ 0.

### Calculating *ℓ*_2,*mut*_

We compute *ℓ*_2,*mut*_ using the distance distribution. Let *D*(*α*) represent the probability that the distance between two adjacent syncmer positions is *α* – 1, and *D_mut_*(*α*) be the same under mutation, then the expressions for *ℓ_mut_* and *ℓ*_2,*mut*_ can be written as:

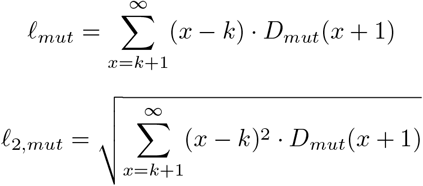

To calculate *D*(*α*) we define the new quantity *F*(*α*) denoting the probability that *only* the first or *only* the last *k*-mer in a window of *α k*-mers is a syncmer, respectively. We refer to these *k*-mers as *K*_1_ and *K_α_* respectively. Note that for *P*(*α*) defined as above, 1 – *P*(*α*) gives the probability that *no k*-mer in an *α*-window is a syncmer.

We compute *F*(*α*) by conditioning on *β* as before. For simplicity we divide the sum over *β* into cases based on the syncmers that are generated by *s_β_*. With some abuse of notation, we let *K_i_* represent the event that *s_β_* generates *K_i_* as a syncmer.

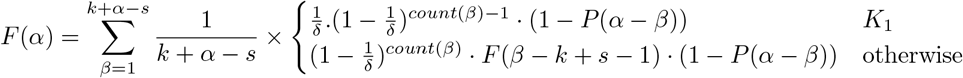

In the first case we have the probability that *K*_1_ is not downsampled, any other syncmer generated by *s_β_* is downsampled, and there are no other syncmers generated to the right of *β*. In the second case we have the probability that any syncmers generated by *s_β_* are downsampled, no syncmers are generated to the right of *β*, and the recursive computation of the probability that the *s*-minimizer of the segment to the left of *β* generates a syncmer at *K*_1_.

Similarly, *D*(*α*) is the probability that in a window of *α k*-mers *only* the first *and* last *k*-mers are syncmers. Then

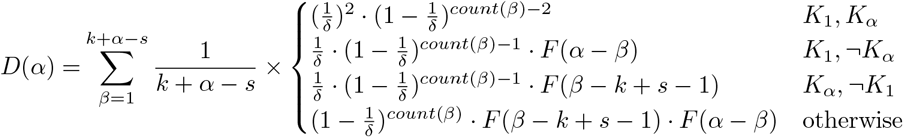

To compute *ℓ*_2,*mut*_ we need the analogous expressions *F_mut_*(*α*) and *D_mut_*(*α*). The expressions for these values are more involved, and left to Supplementary section S2.3. After computing the theoretical *ℓ*_2,*mut*_ according to these expressions, we use it select the best PSS for any compression.

Because of the recursive nature of the theoretical expressions, their computation even to a fixed accuracy is time consuming. In practice, simulating a very long sequence, selecting syncmers, and simulating mutations to determine this metric empirically is less time consuming. Our tests show that the theoretical and empirical results are very close. For example, for 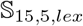 (*i,j*) for 1 ≤ *i* < *j* ≤ 11 and 15% mutations, the average difference was 0.26%. (See Supplementary Table S1 and Supplementary Data File 1, Table 3). We used this simulation method to compute *ℓ*_2,*mut*_ for *k* = 11, 13, 15, 17 and 19, mutation rates 0.05,0.15 and 0.25, and all 2- and 3-parameter schemes. The results are presented in Supplementary Data File 1, Table 2 (note that for 1-parameter schemes the best *ℓ*_2_ and *ℓ* are the same, and thus already known from [9]). *ℓ*_2,*mut*_ values computed using the exact theoretical expressions for some parameter combinations are available in Supplementary Data File 1, Table 3.

**Table 1.** Reference genomes. Basic information about the reference genomes used in our experiments. # scaffolds is the number of individual sequences present in the reference genome fasta file and can include unplaced scaffolds, alternates, etc. Length is the total length (in nt) of all of the scaffolds together, excluding ambiguous bases.

| Dataset | source | Species | # scaffolds | Length |
| --- | --- | --- | --- | --- |
| GRCh | GRCh38 [13] | Human | 639 | 3.111G |
| CHM13X | CHM13 chrX [14] | Human | 1 | 154.3M |
| BAC | PacBio [16] | Microbial | 24 | 59.1M |
| ECK12 | GCF_000005845.2 [15] | *E. coli K-12* | 1 | 4.6M |

**Table 2.** Reads information. The long read datasets used in our experiments. Source names are from Table 1 where relevant. PB=PacBio, ONT=Oxford Nanopore Technologies.

| Dataset | Source | Read type | # reads | Mean length (std) |
| --- | --- | --- | --- | --- |
| pbsim_x | CHM13X | PB simulated | 173891 | 8871.1 (5570.1) |
| pbsim_bac | BAC | PB simulated | 66428 | 8894.2 (5617.4) |
| ns_chm13 | CHM13 | ONT simulated | 1000 | 8722.8 (7030.7) |
| pb_bac | BAC | PB real | 15000 | 9488.3 (5207.2) |
| ont_na12878 | ERR3279003 | ONT real | 10000 | 7131.6 (2348.5) |

**Table 3.** Performance metrics of minimizer and syncmer schemes on real sequences with simulated mutations. Substitutions were introduced in the references at a rate of 15%. The values shown are for the conserved selected *k*-mers. # conserved is the number of *k*-mers selected by a scheme that were conserved under mutation. Best performance is shown in bold.

| Dataset | Scheme | Description | Comp-<br>ression | $\ell$ | $\ell_2$ | $p90$ | $p100$ | # conserved |
| --- | --- | --- | --- | --- | --- | --- | --- | --- |
| ECK12 | $\mathcal{M}_{15,10}$ | minimap minimizer | 76.65 | 0.86 | 13.77 | 211 | 1045 | 60,557 |
| | $\mathbb{S}_{15,5}(3,9)$ | optimal PSS | 62.91 | **0.84** | **12.97** | **182** | **941** | 73,779 |
| | $\mathbb{S}_{15,5}(1,11)$ | closed syncmer | 63.37 | 0.85 | 13.42 | 188 | 1277 | 73,245 |
| | $\mathbb{S}_{15,5}(1,2)$ | “bad PSS” | 63.23 | 0.87 | 14.19 | 195 | 1443 | 73,413 |
| | $\mathcal{M}_{15,19}$ | minimap minimizer | 154.13 | 0.91 | 17.85 | 378 | 1981 | 30,115 |
| | $\mathbb{S}_{15,6}(6)$ | optimal PSS | 116.04 | **0.90** | **16.18** | **303** | **1542** | 40,001 |
| CHM13X | $\mathcal{M}_{15,10}$ | minimap minimizer | 54.29 | 0.81 | 11.70 | 152 | 1219 | 2,841,498 |
| | $\mathbb{S}_{15,5}(3,9)$ | optimal PSS | 45.65 | **0.80** | **11.12** | **132** | **1193** | 3,379,361 |
| | $\mathbb{S}_{15,5}(1,11)$ | closed syncmer | 44.71 | 0.81 | 11.46 | 134 | 1248 | 3,450,241 |
| | $\mathbb{S}_{15,5}(1,2)$ | “bad PSS” | 43.87 | 0.82 | 12.07 | 137 | 1387 | 3,515,931 |
| | $\mathcal{M}_{15,19}$ | minimap minimizer | 107.91 | 0.88 | 15.12 | 270 | 1946 | 1,429,555 |
| | $\mathbb{S}_{15,6}(6)$ | optimal PSS | 83.20 | **0.86** | **13.83** | **219** | **1927** | 1,854,097 |

### Achieving the target compression

A simple extension of the expression for compression of open and closed syncmers yields that the compression of an *n*-parameter PSS is 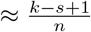, where we assume that *s* is long enough relative to *k* so that the *s*-minimizer is likely to be unique. In Supplementary Data File 1, Table 4 the *ℓ*_2,*mut*_ values for schemes that achieve the same compression either by using more parameters or by downsampling. It is preferable to achieve a specific compression with minimal downsampling. For example, the *ℓ*_2_ of the best 2-parameter scheme with a downsampling rate of 2 is an order of magnitude worse than that of the best 1-parameter scheme that has the same compression without downsampling. Thus, to choose the best PSS for a given target compression, we can choose one with parameters that yield the compression closest to, but below, the desired compression and then downsample to reach the desired compression.

**Table 4.** Runtime and memory. Time (in seconds) and RAM (in GB) needed to index the reference and map the simulated reads by each of the tools. The second and third dataset use the same reference. Syncmer variant parameters were selected to match the minimap2 compression rates as above.

| Task | Method | Scheme | Index<br>time | Index<br>mem | Map<br>time | Map<br>mem |
| --- | --- | --- | --- | --- | --- | --- |
| bacterial reads<br>vs BAC | minimap2 | $\mathcal{M}_{15,10}$ | **3.29** | 0.32 | **11.10** | 3.14 |
| | Syncmer minimap | $\mathcal{S}_{15,5}(3,9)$ | 3.81 | **0.31** | 11.63 | **2.93** |
| ChrX reads<br>vs CHM13X | minimap2 | $\mathcal{M}_{15,10}$ | **7.96** | 1.01 | **65.29** | **5.35** |
| | Syncmer minimap | $\mathcal{S}_{15,5}(3,9)$ | 9.52 | **0.99** | 141.05 | 6.73 |
| | minimap2 | $\mathcal{M}_{15,100}$ | **4.4** | 0.45 | 59.05 | 16.06 |
| | Syncmer minimap | $\mathcal{S}_{15,4}(6)$ | 7.12 | 0.45 | **47.83** | **7.56** |
| bacterial reads<br>vs CHM13X | minimap2 | $\mathcal{M}_{15,10}$ | As | As | **25.61** | **8.26** |
| | Syncmer minimap | $\mathcal{S}_{15,5}(3,9)$ | above | above | 46.59 | 9.28 |

Note that while for 1-parameter PSS the best scheme always has its *s*-minimizer in the middle position as shown by Shaw and Yu [9], for multi-parameter PSS the best scheme may change depending on mutation rate and compression, there is no scheme that is always best in every setting unlike for 1 parameter schemes. The table in Supplementary Data File 1, Table 2 is used to select the best scheme for a given setting.

### Implementing PSS in mappers

We modified the code of minimap2 (v2.22-r1105-dirty) and Winnowmap2 (v2.03) to select our syncmer variants as seeds instead of minimizers. The code for our new syncmer-based mappers is available from https://github.com/Shamir-Lab/syncmer_mapping.

The implementation of the syncmer schemes defined in Definitions and Background is straightforward. Sequences are scanned from left to right, the canonical *k*-mer at each position is identified using a random hash function *h*_1_, and the index of the minimum *s*-mer under another random hash *h*_2_ is determined, and compared against the list of allowed positions of the PSS.

For downsampled schemes, syncmers are selected if their hash value normalized between 0 and 1 is below 1/*δ* where *δ* is the downsampling rate. Note that a different hash function than *h*_1_ must be used to ensure random downsampling. Windowed schemes are integrated into the minimizer selection scheme of the mappers except that syncmers are selected in each window first. If no syncmer is present, then the minimizer is selected.

Pseudocode describing these implementations is presented in Algorithm 1 for regular PSS and in Algorithm S1 in Supplementary section S3 for windowed PSS. Additional implementation and optimization details are presented in Supplementary section S4.

#### Algorithm 1 Syncmer selection (regular, downsampled)

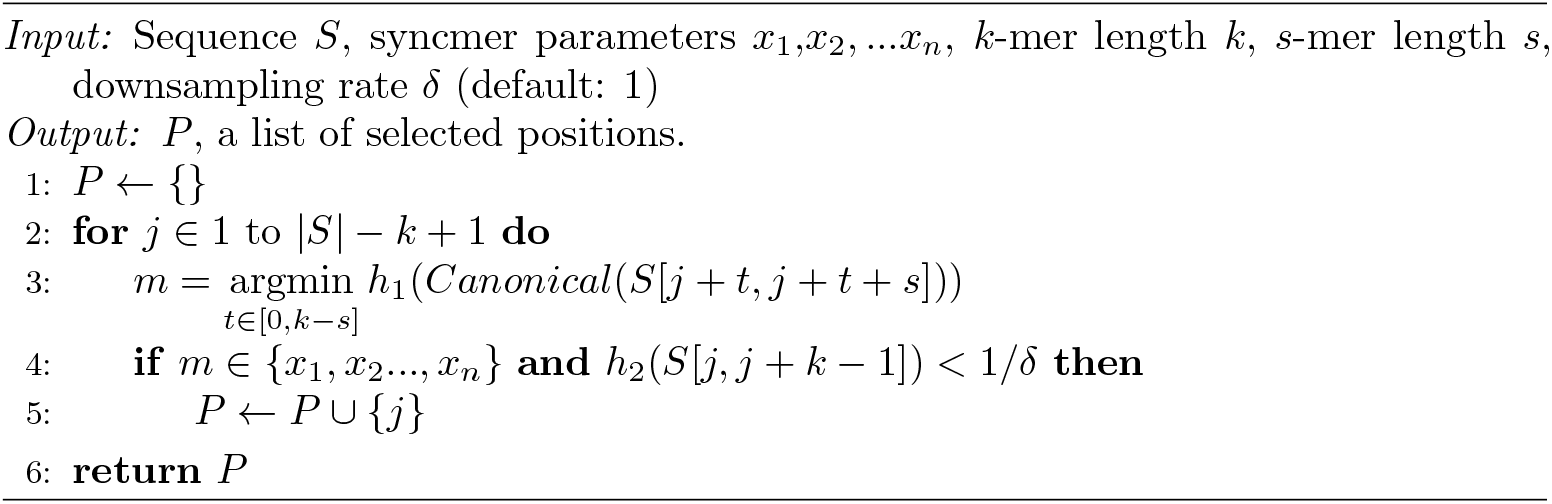

## Results

We first evaluated different PSS on real genomes to demonstrate their properties compared to the theoretical analysis presented above. We then compared PSS-based mapping to the original minimizer-based versions of minimap2 and Winnowmap2 on simulated and real read data.

The reference sequences used for these experiments were: human genome GRCh38.p13 [13], human chromosome X from CHM13 (v1.0) [14], *E. coli* K12 [15], and a set of microbial genomes that we will call BAC, containing assemblies of 15 microbes for which PacBio long read data is available [16] (three of the microbes were used in [9], see Supplementary Section S6 for more details). Information about the sequences is presented in Table 1.

We simulated PacBio and ONT reads from the human genome and from BAC with a depth of 10 using PBSIM [17] and NanoSim [18]. Details of simulation parameters are found in Supplementary section S5.

For tests on real read datasets we selected a random set of 10K ONT reads of the NA12878 cell line with read length capped at 10kb (SRA accession ERR3279003), and 1K PacBio reads for each of the BAC microbes [16]. Details are available in Table 2.

### Performance of parameterized syncmer schemes

Our theoretical analysis of PSS properties above relies on a number of assumptions. Specifically, it assumes uniform iid sequences and mutations, allows substitutions only, and treats the sequence as a single forward strand. We therefore examined the properties of PSS on real genomes where these assumptions do not necessarily apply, and compared them to minimizer schemes.

We used *k* = 15 and selected the best syncmer schemes (based on *ℓ*_2,*mut*_) with theoretical compression 5.5 and 10 (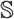(3, 9) and 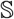(6), respectively). The default minimizer scheme of minimap2 uses *k* = 15 and *w* = 10 yielding the theoretical compression of 5.5. A theoretical compression of 10 is achieved by minimap2 with *k* =15 and *w* = 19. For compression 5.5 we also included in the comparison closed syncmers (i.e. 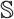(1,11)) and a syncmer scheme that should perform poorly according to the theoretical analysis (“bad PSS”, 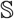(1, 2)). We compared the schemes on both unmutated sequences and on sequences with iid substitutions simulated at a rate of 15%. Since conservation is defined for index-preserving mutations, indels were not simulated (sequencing errors were included in all subsequent simulations in the following sections).

We tested the schemes on the ECK12 and CHM13X sequences, with and without mutations. On *unmutated* reference sequences, minimizers outperformed syncmers, with much lower *ℓ*_2_ and *p* 100 values for schemes with the same compression (Supplementary section S7, Table S2). The theoretically best PSS outperformed the closed syncmer scheme and the “bad PSS”. In contrast, under mutation, the advantage of syncmers is clear (Table 3): PSS had better performance in all metrics, with the theoretically best PSS performing better than minimizers and closed syncmers. While syncmers had significantly more conserved positions than minimizers, the “bad PSS” 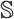(1, 2) had a worse distribution of selected positions and thus poorer *ℓ* and *ℓ*_2_than minimizers.

We also wished to test the impact of using canonical *k*-mers on the distance distribution between selected positions. Fig S2 shows the distance distributions for syncmers selected only using forward strand *k*-mers and using canonical *k*-mers. We conclude that while the theory is limited to single-stranded sequences it shows trends that hold for canonical *k*-mers. Further details can be found in Supplementary section S8.

### The fraction of unmapped reads

We mapped reads using minimap2 and Winnowmap2 with 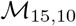 (low compression), 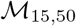 (medium), and 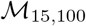 (high) on four datasets. For each dataset, syncmer-minimap and syncmer-winnowmap parameters yielding the best theoretical *ℓ*_2,*mut*_ for the same compression achieved by minimap2 were selected. This resulted in 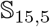(3,9) matching the low compression, and 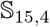(6) matching the medium and high compression. The downsampling rate was manually selected to closely match the real compression of the corresponding minimizer scheme. The exact compression and downsampling rates are given in Supplementary Data File 1, Table 5.

Fig 4 (top) shows the percentage of unmapped reads of the mappers for simulated PacBio and ONT reads mapped to the human reference genome. See Supplementary Fig S3 for additional results, including windowed mappers. Syncmer variants performed essentially the same or better than the original mappers in all cases, with the largest advantage at high compression. All mappers did much better on the PacBio reads than on ONT reads, which have a higher proportion of deletions and substitutions. The jump in the fraction of unmapped reads between medium and high compression may indicate that in order to overcome the large fraction of non-conserved seeds, existing mappers need to use a lower compression with many redundant seeds.

**Fig 4.**
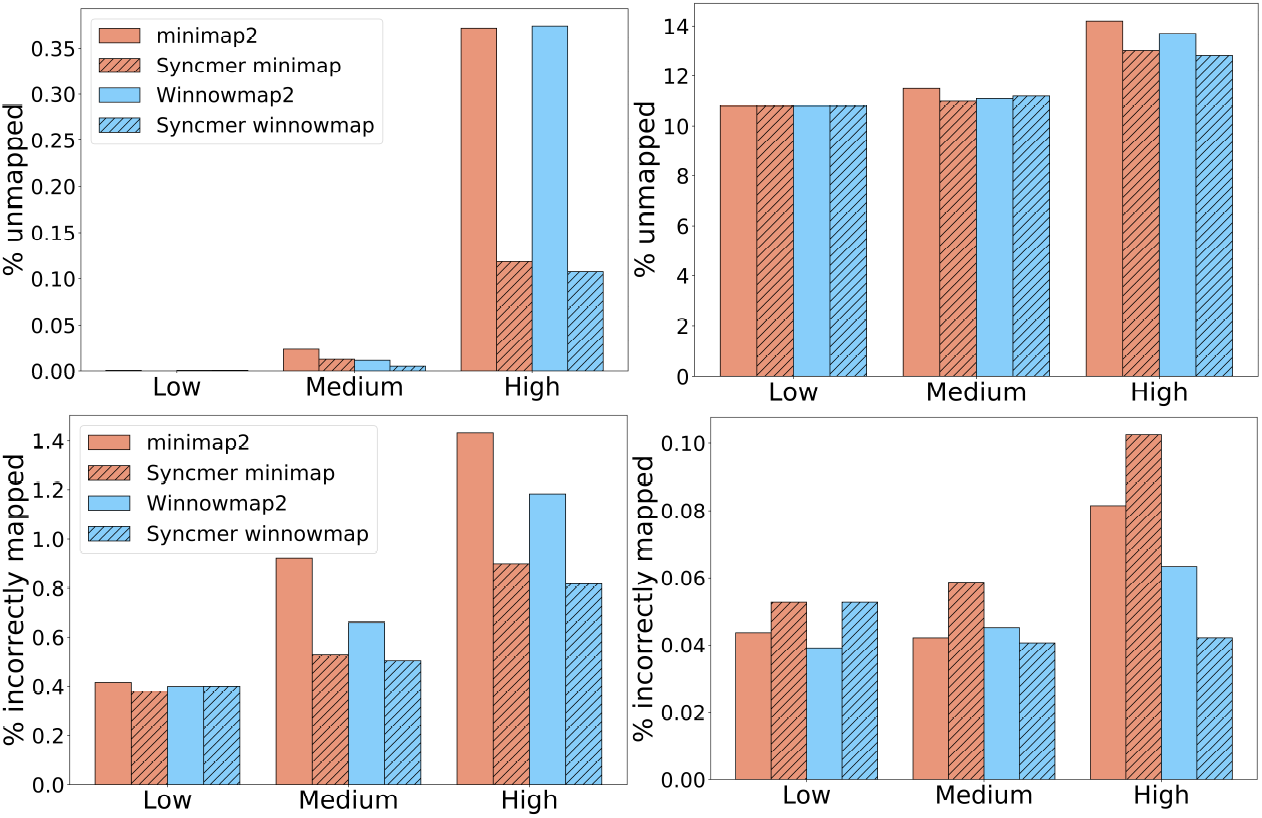
The percentage of unmapped and incorrectly mapped reads - simulated data. **Top:** Percent unmapped for low, medium and high compression. Left: PacBio reads simulated from the CHM13X sequence mapped against ChrX sequences from GRCh38; Right: 1000 ONT reads simulated from CHM13 mapped against GRCh38. **Bottom:** The percentage of incorrectly mapped reads for low, medium and high compression. Left: on PacBio reads simulated from the CHM13 ChrX sequence mapped against CHM13X; Right: PacBio reads simulated from the 15 bacterial species in BAC pooled together and mapped against the union of their references.

We compared the performance of all mappers on real data (Table 2) across a range of compression values. The ONT reads were mapped against the human reference GRCh and the PacBio bacterial reads were mapped against the BAC reference. For the original minimap2 and Winnowmap2 different values of compression were achieved by varying *w*. For the syncmer variants, schemes were selected with the best *ℓ*_2,*mut*_ according to the theoretical analysis and then downsampled as discussed above (Achieving the target compression). Results are shown in Fig 5. The syncmer variants had consistently higher percentage of mapped reads than the original minimizer-based mappers, with syncmer-winnowmap performing the best across the larger part of the compression range. For high compression, the minimizers had 20-40% more unmapped reads than the syncmers. At low compression rates of 5.5 –11, minimizers had 2-15% more unmapped reads than syncmers. Full results and scheme parameters are given in Supplementary Data File 1, Table 6.

**Fig 5.**
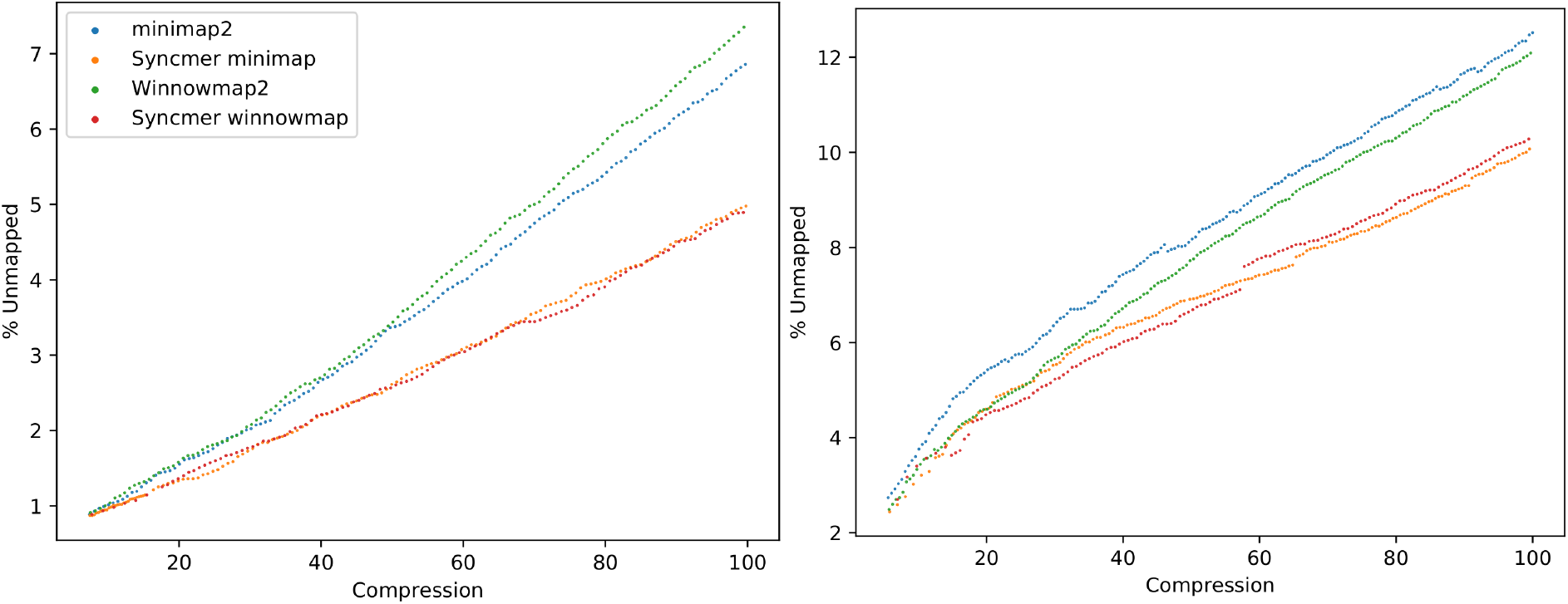
Percentage of unmapped reads – real datasets. Percentage is shown as a function of compression rate, PSS parameters were chosen to achieve the desired compression with lowest *ℓ*_2,*mut*_. **Left:** Pooled PacBio bacterial reads mapped against BAC. **Right:**ONT human cell-line reads mapped against GRCh38.

To compare the performance of PSS to syncmers, we mapped the PacBio bacterial reads against BAC using syncmer-minimap with 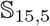(3,9), the theoretically best 2-parameter scheme, and with closed syncmers (equivalent to 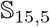(1,11)) across a range of compression values. The results are shown in Fig S4. The PSS selected by our analysis had consistently fewer unmapped reads than closed syncmers. Note that 1-parameter PSS and open syncmers are equivalent and the best scheme always has its *s*-minimizer in the middle position as discussed above and by Shaw and Yu [9].

### Mapping correctness

We evaluated the mapping correctness for PacBio simulated reads as done in [1] (see Supplementary section S5 for details). The percentage of incorrectly mapped reads simulated from CHM13X and the BAC genomes is shown in Fig 4 (bottom). Winnowmap was consistently better than minimap, and the syncmer variants of Winnowmap performed best at medium and high compression.

Although we cannot evaluate the mapping correctness on the real datasets, the mapping quality scores reported by minimap2 can be used to compare the different mappers. On the real datasets, reads mapped by syncmer-minimap but not by minimap2 generally had higher mapping quality than those mapped by minimap2 and not syncmer-minimap. For example, for the human cell line ONT reads, comparing minimap2 with 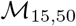 to syncmer-minimap, the 39 minimap-only reads had average mapping quality 31.4 (median 27), while the 94 syncmer-minimap-only reads had an average quality score of 38.7 (median 42.5). Full results for different compression rates are presented in Supplementary Data File 1, Table 7.

### Impact of sequence identity level

We examined the impact of the level of identity between the sequenced reads and the reference to which they are aligned. Differences between the sequences can be due to sequencing errors, mutations in the sequenced organism, or differences between sequenced and reference strains. We simulated 1000 PacBio reads from CHM13X at percent identity 65%, 75%, 87% and 95%. The results are shown in Fig 6 and S7. For minimap2 and Winnowmap2 we used 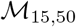, and in the syncmer variants we used 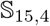(6) with the other parameters selected as above to match the compression of minimap2.

**Fig 6.**
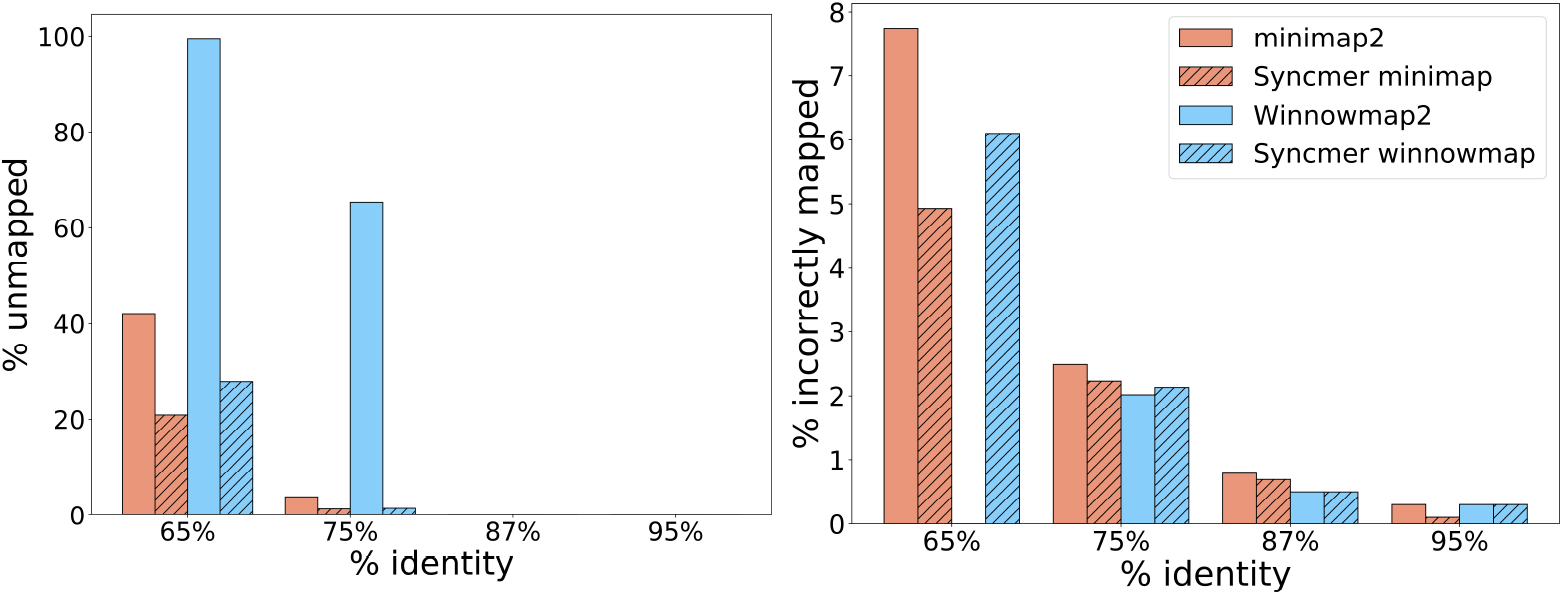
Impact of percent sequence identity on mapping quality. We varied the mutation rate of 1000 PacBio simulated reads from CHM13X. The figures present the % unmapped and incorrectly mapped by each method. **Left:** % unmapped reads. **Right:** % of the mapped reads that were incorrectly mapped.

The syncmer variants outperformed the original tools in terms of fraction of reads mapped, with larger gains as percent identity decreases. All tools performed very well at higher percent identity, indicating that more than enough seeds were selected and conserved to adequately map all reads (and thus perhaps compression could be increased). Winnowmap2 performed noticeably worse at lower percent identity, leaving almost all reads unmapped at 65% identity. Syncmer-minimap outperformed minimap2 on the fraction of correctly mapped reads in all cases. Winnowmap2 correctly mapped a larger fraction of the mapped reads at 75% identity, but mapped only 35% of the reads, compared to ≥ 95% for the other variants. At 95% identity the syncmer variants had fewer incorrectly mapped reads. While very low percent identity may be unrealistic in some cases, these results highlight the impact of the increased conservation of syncmers.

### Performance of windowed syncmer schemes

Windowed schemes combine syncmers and minimizers, complementing the syncmer scheme to provide a window guarantee. Supplementary section S10 presents the results of the experiments on windowed PSS. Although windowed schemes perform better than the unwindowed on some metrics (compare Tables S4, 3), in practice the windowed variants of our syncmer mappers were similar or worse than the variants without windowing for the same compression in most cases (Figures S5-S8).

### Runtime and memory

We compared the runtime and memory usage of the six tested mappers on a number of datasets. All experiments were performed on a 44-core, 2.2 GHz server with 792 GB of RAM, using 50 threads. Peak RSS (in GB) and real time (in seconds) as measured by the tools are reported.

Table 4 compares the separate performance of indexing and mapping on simulated PacBio and ONT reads from bacteria and human. Winnowmap was not compared as it does not allow for separate indexing and mapping, a disadvantage when many read sets will be mapped to the same reference. For syncmer minimap variants the same parameters matched to the minimizers as above were used. At low compression minimap2 had better runtimes for both indexing and mapping, and memory usage was similar between the tools. At high compression syncmer-minimap had longer indexing time but lower mapping time and required less than half the memory. This is in addition to having only 1/3 as many unmapped reads (Fig S3A).

We also compared the runtime and memory of all the runs for different compression rates shown in Fig 5. Results are shown in Figures 7. Note that the results here are for indexing and mapping together. minimap2 was consistently the fastest, followed by syncmer-minimap, which took 50-100% longer. Interestingly, the two datasets show opposite trends in memory usage (Fig 7, bottom). This is because the bacterial reference genomes are relatively short, and thus the memory bottleneck is in the mapping stage, while for the human reference genome the memory bottleneck is in the indexing stage. Increasing compression lowers index size but results in longer alignments between anchors, requiring more memory in the mapping phase. Thus, when indexing is the bottleneck, increasing compression reduces memory, while when mapping is the bottleneck it increases memory. Winnowmap2 and its variants used less memory in the mapping phase while minimap2 and its variants used less memory in the indexing phase. In the case that indexing was the bottleneck, the syncmer variants required lower memory usage than the original mappers across most of the range of compression rates (Fig 7, bottom right).

**Fig 7.**
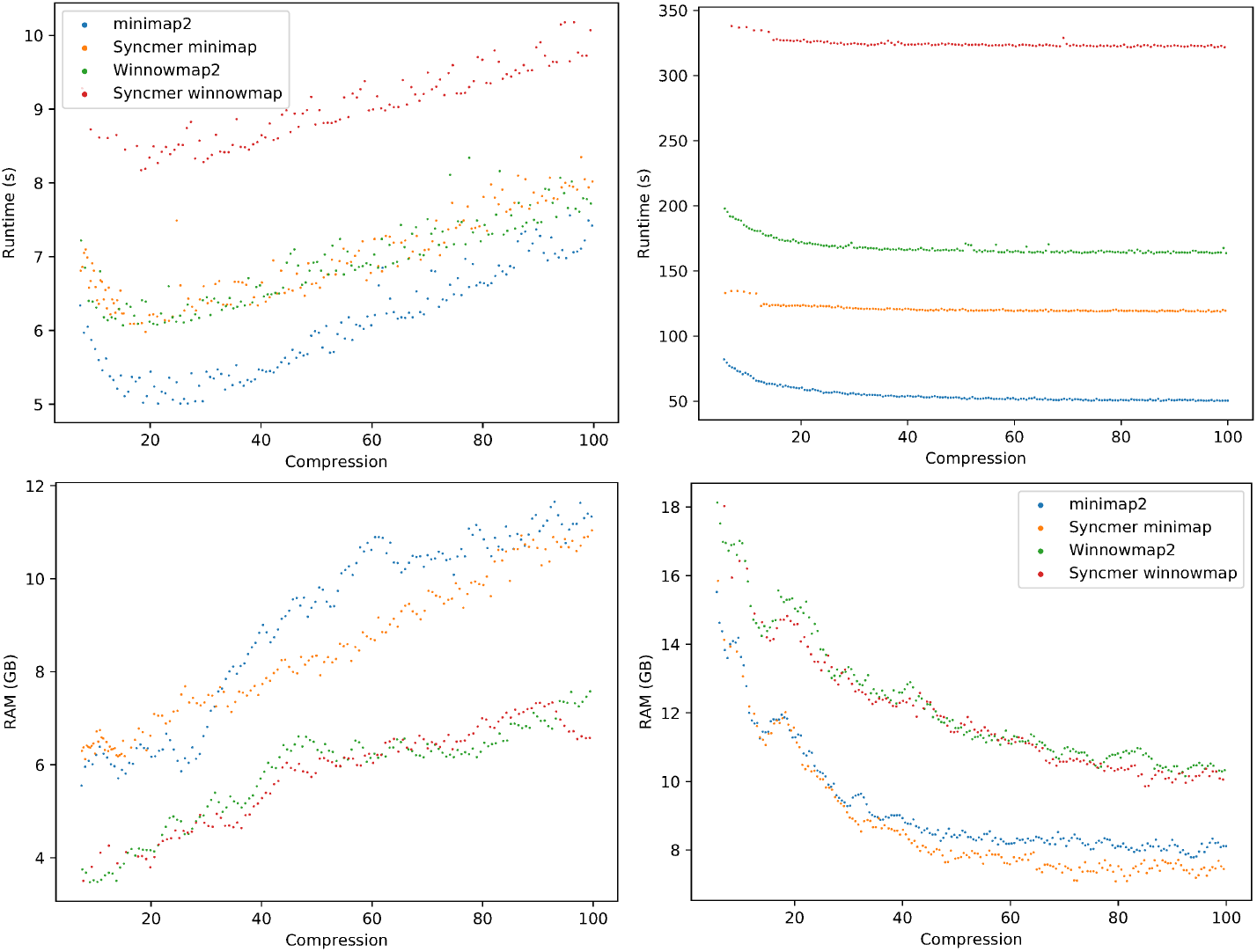
Memory usage and runtime vs. compression – real data. **Top:** Runtime in seconds to index the reference and map reads by each method. **Bottom:** Peak RAM usage in GB to index the reference and map reads. **Left:** PacBio bacterial reads. **Right:** ONT human cell-line reads.

## Discussion

In this study we generalized the notion of syncmers to PSS and derived their theoretical properties. We incorporated PSS into the long read mappers minimap2 and Winnowmap2. Our syncmer mappers outperformed minimap2 and Winnowmap2, by mapping more reads and correctly mapping a higher fraction of those mapped across a range of different compression values for multiple real and simulated datasets.

As our results show, the advantage of using syncmers is most marked at high compression and high error rates, as is expected due to their higher conservation. Yet the advantage is already manifest at the lower compression rates commonly used by existing mappers. For large genomes, such as the human genome, using the higher compression enabled by syncmers also leads to lower RAM usage. Syncmer-minimap is slower than the highly optimized minimap2, taking 50-100% longer to map reads, but it is faster than Winnowmap2. Future work should focus on lowering the runtime by optimizing the syncmer mapping implementation.

The well-developed minimap2 and Winnowmap2 software tools have a variety of internal parameters, and by adjusting them one may be able to achieve some of the performance advantage of PSS. The approach we propose here is more principled and avoids the need to guess or grid search across parameters in order to get the best mapping performance. Increased performance achieved in such a manner could likely improve the syncmer mappers as well, as improving the choice of alignment seeds is orthogonal to many of the other algorithmic details of read mapping.

There are a number of issues and questions that this work leaves open, particularly in the theoretical analysis. First, the analysis of windowed schemes and downsampled schemes under mutation remains to be completed. Second, an expression for *ℓ*_2_ for minimizer schemes could also be obtained. Third, can the theory be expanded to canonical *k*-mers? Fourth, it would be helpful to obtain more robust definitions of conservation and *ℓ*_2_ that do not depend on preserving indices between sequences, thereby allowing indels to be included in the theoretical analysis. Finally, what is the ideal metric for evaluating the performance of schemes? While we argue that *ℓ*_2_ is preferable to *ℓ*, other new metrics may capture mapping performance even more accurately.

Another possible avenue to explore is in the definition of the selection scheme itself. Is it possible to select *k*-mers in a biased way in order to increase the compression but still retain the beneficial distance distribution of syncmer schemes? Or could a sequence-specific set of *k*-mers be determined efficiently for any desired compression rate? The quest for a “best” selection scheme is not over.

## Acknowledgements

We thank members of the Shamir Lab for their helpful comments.

## Supplementary information for

### S1 Definitions of downsampled and windowed syncmer schemes

#### Downsampled syncmers

Given a uniformly random hash function *h*: Σ^k^ → [0, *H*], for a given string *S*, *downsampling* selects syncmers only from the set of |Σ|^k^/*δ k*-mers with the lowest hash values.

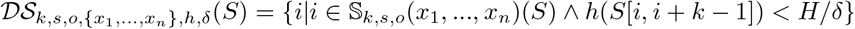

We call *δ* the *downsampling rate*.

#### Windowed syncmers

Windowed syncmers fill in gaps using a minimizer scheme, thus providing a window guarantee. For clarity in the definition below let 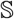(*S*) represent 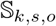(*x*_1_,…, *x_n_*)(*S*).

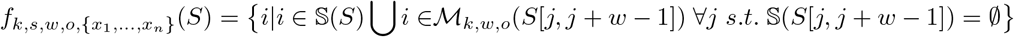

Letting 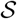 represent all *k*-mers that can be syncmers in 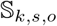(*x*_1_,…, *x_n_*), an equivalent definition would be: 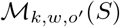 where *o*’ is defined such that 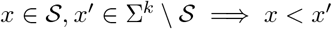.

### S2 Analysis of parameterized syncmer schemes

#### S2.1 Recursive expressions for conservation of PSS

Consider a window of *α* consecutive *k*-mers. We assume randomly distributed sequence throughout. Let *s_β_* be the *s*-minimizer in the window, at position *β*. Then if *t* is a parameter of the syncmer scheme, *s_β_* generates a syncmer if it is not in the first *t* – 1 or last *k* – *t* positions in the window. If *β* is not in a position where it generates a syncmer, we recursively check to the left or right of *β* to see if a syncmer is generated by the *s*-minimizer of that region. See Fig S1 for an example.

For a 1-parameter scheme *f* with *k*-mer length *k*, *s*-minimizer length *s*, and parameter *t* let *P* (*α*) be the probability of selecting at least one syncmer in a window of α adjacent *k*-mers. Then, assuming a uniformly random hash over the *s*-mers, and conditioning on the position of the *s*-minimizer of the *α*-window, *β*:

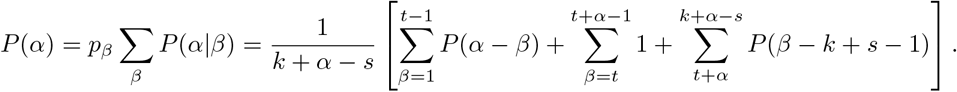

The probability of any of the *k* + *α* – *s* starting positions being the *s*-minimizer is denoted as *p_β_* and assumed to be uniform. If *β* is in the first *t* – 1 or last *k* – *t* starting positions (red regions in Fig S1A), then a syncmer may be generated by the remaining *α* – *β* positions to the right or *β* – *k* + *s* – 1 positions to the left, respectively. Note we define 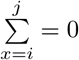 when *i* > *j* and *P*(*x*) = 0 when *x* ≤ 0.

When downsampling syncmers, there is a probability of 1/*δ* that an *s*-minimizer in the syncmer generating region (i.e. the green region in Fig S1A) really generates a syncmer. If it does not, then the left and right regions are considered recursively, yielding the following expression, where we simplify notation by letting *P*(*α* – *β*) = *P_R_* and *P*(*β* – *k* + *s* – 1) = *P_L_*:

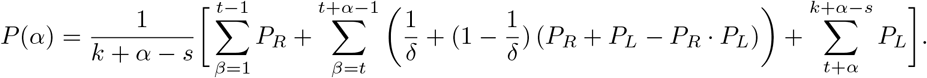

**Fig S1.**
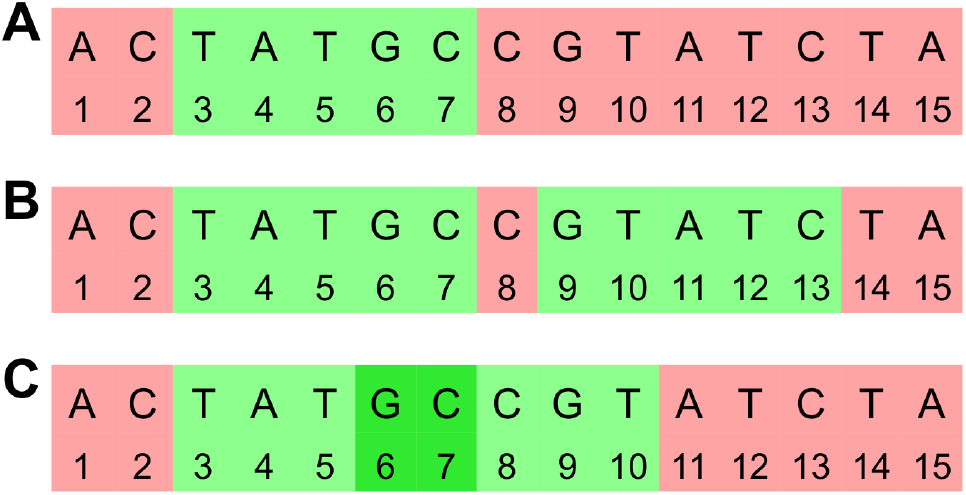
Illustration of *s*-minimizers generating syncmers. A window of *α* = 5 consecutive 11-mers. **A**: When *s* = 5 and *t* = 3, then the *s*-minimizer of the entire window generates a syncmer when its starting index is in the green region. If the *s*-minimizer is in one of the red regions then a syncmer may be generated by the *s*-minimizer of the remaining part of the window. For a two parameter scheme the *s*-minimizer creates two syncmer generating regions that may be disjoint (**B**) if *s* > *t*_2_ – *t*_1_ or overlapping (**C**) if *s* < *t*_2_ – *t*_1_. In this example, *t*_1_ = 3 and *t*_2_ = 9 in **B** and *t*_2_ = 6 in **C**.

In the case of 2-parameter schemes, two syncmers are generated by *s_β_* in regions that will overlap if the parameters *t*_1_ and *t*_2_ are within *s* of each other, and will be disjoint otherwise (see Fig S1B,C). Combining these two cases into a single recursive expression yields:

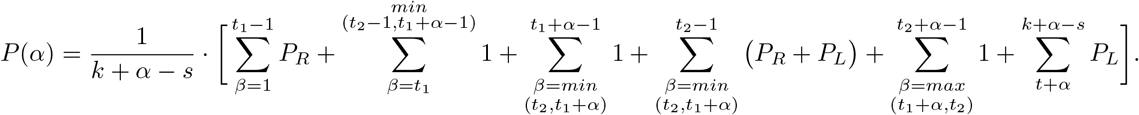

When downsampling is used then the 1 in the second and fifth sums is replaced by 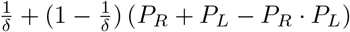 as in the one parameter case. The third sum expresses the overlapped region where either parameter creates a syncmer, when it exists. When *both* generated syncmers are downsampled then the left and right sides are recursively checked, thus the 1 is replaced by 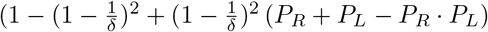.

This expression can be greatly simplified by introducing the notation *count*(*β*) that represents the number of syncmers generated by the *s*-minimizer *s_β_*. For example, *count*(*β*) = 0 in the red region of Fig S1 and *count*(*β*) = 2 in the overlapped region when *β* = 6 or 7 in Fig S1C. The general expression for *P*(*α*) for *any* PSS is:

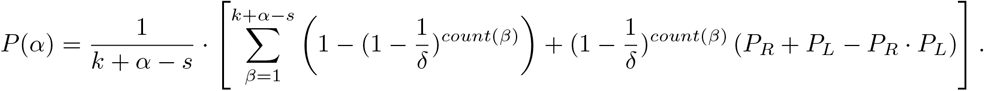

Note that this definition relies on the definition *P*(*x*) = 0, *x* ≤ 0 to include the correct terms for the correct values of *β*.

The value of *P*(*α*) can thus be computed efficiently for any PSS and used to calculate the conservation using the formula from [1].

#### S2.2 Calculating the distance distribution

For a given scheme, the distribution of distances between adjacent syncmer positions is specified by *Pr*(*d* = *x*), the probability that the distance *d* is *x*. To calculate this probability, we define the new quantities *F*(*α*) and *L*(*α*) denoting the probability that *only* the first or *only* the last *k*-mer in a window of *α k*-mers is a syncmer, respectively. We refer to these *k*-mers as *K*_1_ and *K_α_* respectively. Note that due to symmetry *F*(*α*) = *L*(*α*). Note also that 1 – *P*(*α*) gives the probability that *no k*-mer in an α-window is a syncmer.

We compute *F*(*α*) by conditioning on *β* as before. For simplicity we divide the sum over *β* into cases based on the syncmers that are generated by *s_β_* rather than breaking up the sum across different values of *β*. With some abuse of notation, we let *K_i_* represent the event the that *s_β_* generates *K_i_* as a syncmer.

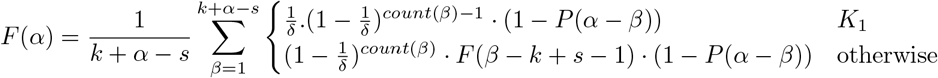

Similarly, define *D*(*α*) to be the probability that in a window of *α k*-mers *only* the first *and* last *k*-mers are syncmers. Then

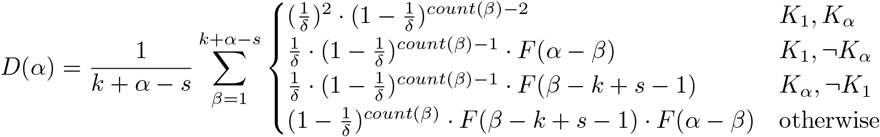

#### S2.3 Calculating *ℓ*_2,*mut*_

To compute the desired metric *ℓ*_2,*mut*_ we must calculate the metrics from the previous section but only with *conserved* syncmers. We add the subscript ‘*mut*’ to a value to indicate that only conserved syncmers are considered. The impact of mutations is similar to that of downsampling shown in the previous section, except that when a syncmer is lost due to mutation, the surrounding *k*-mers are also lost. In this case we consider no downsampling to make the expressions simpler.

Let Ω_*β*_ be the set of syncmers generated by *s_β_*, and *ω_β_i__* be the members of this set. Note that |Ω_*β*_| = *count*(*β*). Then,

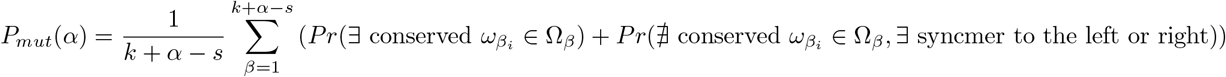

For convenience we call the first probability *P_conserved_* and the second *P_recurse_*.

*P_conserved_* is computed using the inclusion-exclusion principle:

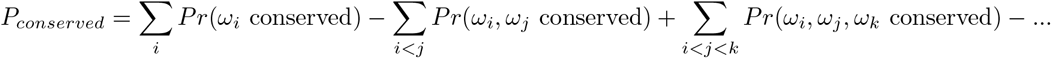

Every term in this series is calculated as (1 – *θ*)^*count Bases*^ where *θ* is the mutation rate and *count Bases* counts the number of bases covered by the conserved *k*-mers (i.e. if two conserved syncmers overlap, the shared bases are counted only once).

*P_recwrse_* is more complicated. We again sum over all values of *β*. When Ω_*β*_ is empty (e.g. *β* is in the red region), then the recursion is similar to the case without mutation. Otherwise, all of the syncmers are lost due to mutation, and we additionally sum over the possible positions of the first and last points of mutation in Ω*β*, named *f* and *l*, respectively.

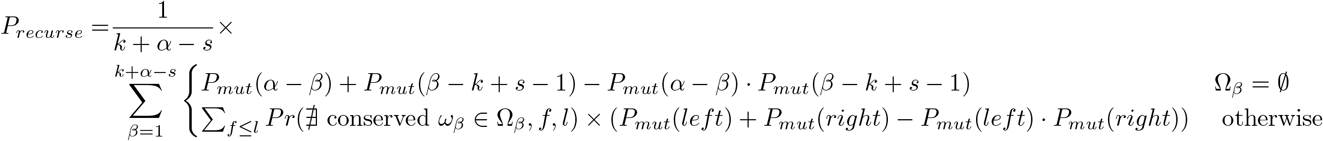

Here *left* = *max*(0, *min*(*f* – *k*, *β* – *k* + *s* – 1)) and *right* = *max*(0, *min*(*α* – *l*, *α*– *β*)). We expand the joint probability as

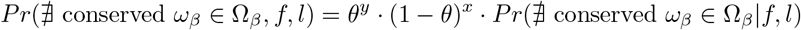

where *y* is 1 if *l* = *f* and 2 otherwise, and *x* is the number of unmutated bases that is fixed by the given values of *f* and *l*. The conditional probability is written as

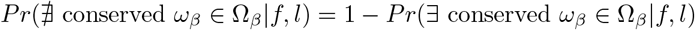

and is computed using the inclusion-exclusion principle as above.

*F*(*α*) and *D*(*α*) are extended to the case of mutation similarly:

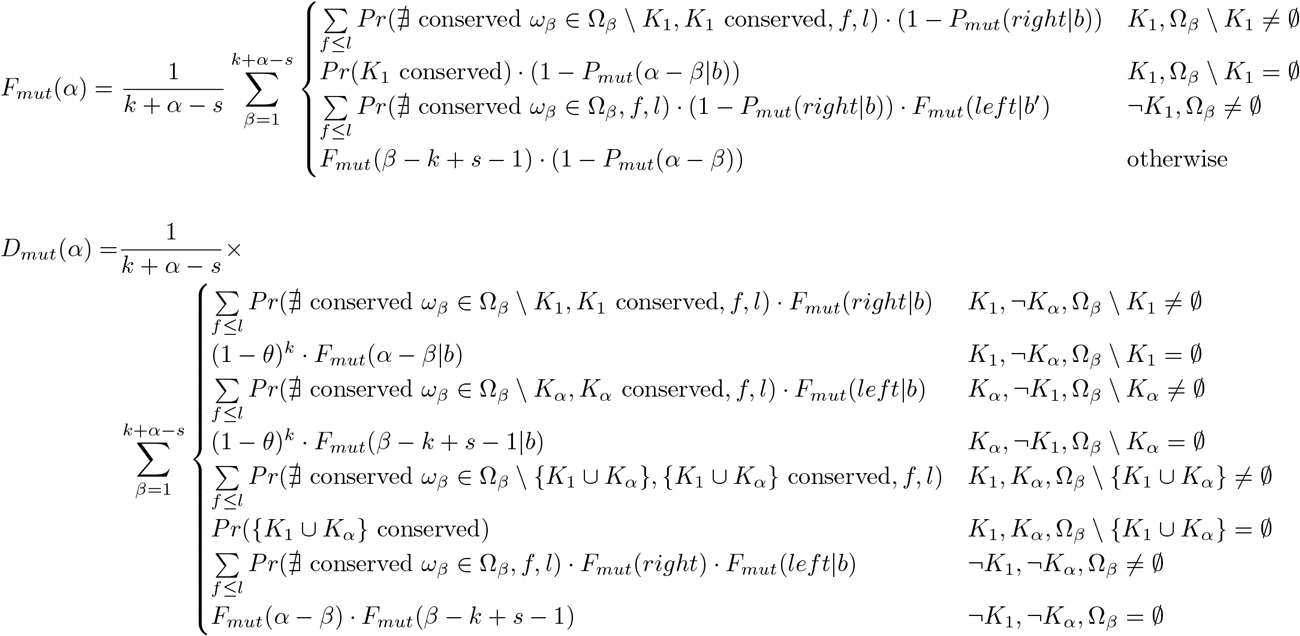

**Table S1.**
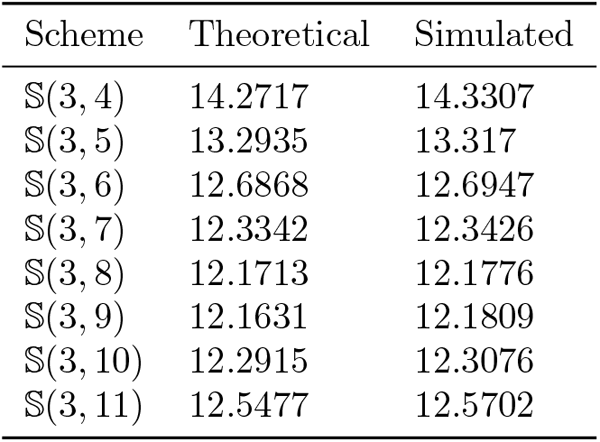
Comparison of theoretical and simulated values of *ℓ*_2,*mut*_. Values of *ℓ*_2,*mut*_ for a selection of schemes with *k* = 15, *s* = 5, *θ* = 0.15. Theoretical values were computed using 1200 terms. Simulated values were found on a simulated sequence of length 1,000,000.

Here we again divide into cases depending on whether there are syncmers that can be lost. We have also used recursive expressions that are similar to the above except we are given that *b* bases to the left or right of the defined region are conserved. These are calculated using similar techniques as above.

Finally, we can use these expressions to compute:

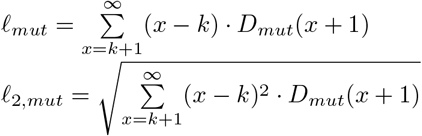

Note that, unlike *P*(*α*), which can be computed efficiently, the computation of these metrics includes an infinite sum. The sum can be truncated at an appropriate distance *x*, however there are still many more terms than in the computation of the conservation. In practice, simulating a very long sequence, selecting syncmers, and simulating mutations to determine these metrics empirically is much less time consuming and yields results that are very close to the true values. We used this simulation method to compute *ℓ*_2,*mut*_ for *k* = 11, 13, 15, 17 and 19, mutation rates 0.05, 0.15 and 0.25, and all 2- and 3-parameter schemes Results are presented in Supplementary Data File 1, Table 2 (note that for 1-parameter schemes the best *ℓ*_2_ and *ℓ* are the same, and thus already known from [1]). The exact theoretical expressions for some parameter combinations are available in Supplementary Data File 1, Table 3. A comparison of the values computed theoretically and by simulation for the 2-parameter schemes 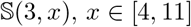 with *k* = 15, *s* = 5, 15% mutation rate can be seen in Table S1.

### S3 Windowed PSS implementation

Algorithm S1 describes the implementation of windowed PSS.

### S4 Syncmer based mapping implementations

Modifications to the mappers were minimal. Only the code that selects the *k*-mers to use as seeds to index the reference sequence and as anchors from the query reads was modified. Here we describe implementation details and optimizations in the code that differ from the high-level descriptions in Algorithm 1 and S1.

#### Algorithm S1 Windowed syncmer selection

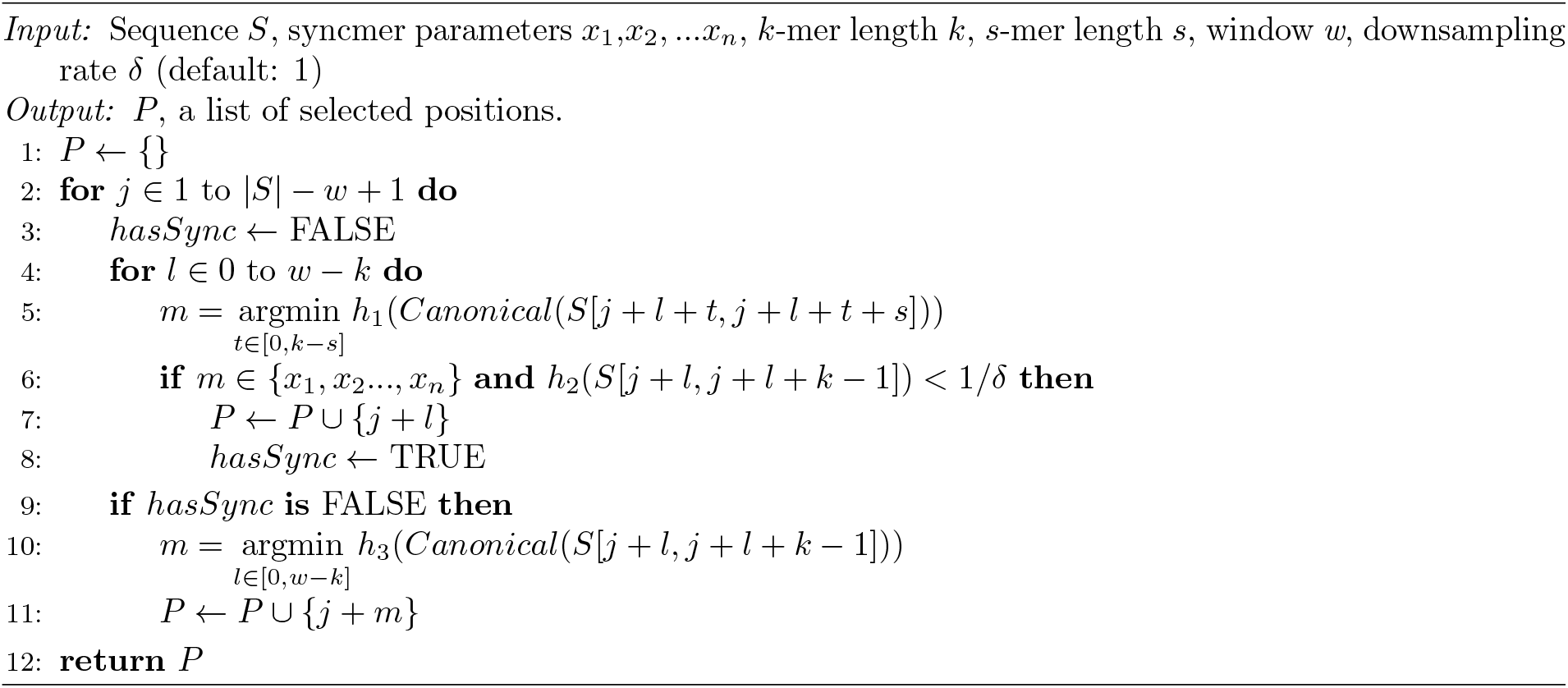

In minimap2 the most frequent minimizers (0.02% by default) are dropped to reduce spurious matches and lower the runtime and memory usage. We also drop the most frequent selected *k*-mers as the last stage of all minimap syncmer variants for consistency. In Winnowmap, the most frequent *k*-mers (also 0.02% by default) are re-weighted in the minimizer order so they are less likely to be selected as minimizers. In the syncmer-winnowmap variant, we do not consider *k*-mer weighting, and thus we simply drop these *k*-mers if they are selected. However, in the *windowed* syncmer-winnowmap variant we do re-weight the frequent *k*-mers before selecting minimizers in empty windows.

We use several different hashes in our syncmer variants of the mappers: *h_can_* to select canonical *k*-mers, *h_s_* to select *s*-minimizers, *h_min_* to select minimizers for windowed variants and *h_down_* for downsampling. We require that *h_can_* ≠ *h_down_* to maintain random downsampling. In minimap syncmer variants we use hash64 from minimap2 for *h_min_* and a variant of MurmurHash2 that ensures *murmur*2(0) ≠ 0 to ensure randomness for the other hashes. Thus *h_min_* = – *hash*64/UINT64_MAX, *h_s_*(*x*) = *murmur*2(*x*), *h_down_*(*x*) = *murmur*2(*x*), and *h_can_*(*x*) = *murmur*2(*x* << 1 + 5) to ensure that it has a different value than *h_down_*. For winnowmap variants we use *h_can_*(*x*) = *lexicographic*(*x*) as this is what is used by the *k*-mer counter Meryl, *h_min_(x)* = –(*murmur*2(*x*)/UINT64_MAX)^8^ in the case that the minimizer is one of the most frequent and – *murmur*2(*x*)/UINT64_MAX otherwise. The other hashes are as in the minimap variants.

In all windowed variants, downsampling occurs before filling in empty windows with minimizers.

ONT reads were mapped using the `map-ont` option in all mappers, while PacBio reads were mapped using the `map-pb` option (`map-pb-clr` in Winnowmap and variants). The latter uses homopolymer compression (HPC) and thus has a real compression (on the non-HPC sequence) that is above the theoretical one.

### S5 Simulation details

PacBio reads were simulated using PBSim [2] with the CLR model and the following parameter settings: depth 10, mean length 9000, length std 6000, minimum length 100, maximum length 40000, mean accuracy 0.87, accuracy std 0.02, minimum accuracy 0.85, maximum accuracy 1, and difference ratio 10:48:19. Error rates and read lengths roughly matched to the statistics observed in a recent benchmark of long read correction methods [3] unless otherwise indicated.

ONT reads were simulated using NanoSim [4] with default parameters and the human pre-trained model for Guppy base calls.

To evaluate mapping correctness for the PacBio simulated data we used the mapeval utility of paftools packaged with minimap2. In this tool, reads are considered correctly mapped if the overlap between the read alignment and the true read location is ≥ 10% of the combined length of the true read and aligned read interval. This criterion was also used in [5].

### S6 Bacterial species used

We chose a single representative assembly of each strain from [6] with the fewest, longest and most highly covered contigs and concatenated all references into a single fasta file. Reads from the same samples were downloaded. Assemblies and reads from following samples were used:

- bc1019, Bacillus cereus 971 (ATCC 14579)
- bc1059, Bacillus subtilis W23
- bc1101, Burkholderia cepacia (ATCC 25416)
- bc1102, Enterococcus faecalis OG1RF (ATCC 47077D-5)
- bc1111, Escherichia coli K12
- bc1087, Escherichia coli W (ATCC 9637)
- bc1018, Helicobacter pylori J99 (ATCC 700824)
- bc1077, Klebsiella pneumoniae (ATCC BAA-2146)
- bc1082, Listeria monocytogenes (ATCC 19117)
- bc1043, Methanocorpusculum labreanum Z (ATCC 43576)
- bc1047, Neisseria meningitidis FAM18 (ATCC 700532)
- bc1054, Rhodopseudomonas palustris
- bc1119, Staphylococcus aureus HPV (ATCC BAA-44)
- bc1079, Staphylococcus aureus subsp. aureus (ATCC 25923)
- bc1052, Treponema denticola A (ATCC 35405)

### S7 Performance of minimizers and PSS on real genome sequences without mutations

The metrics measured on real genomes without mutations are shown in Table S2.

### S8 Distribution of distances between canonical syncmers

To test the impact of using canonical *k*-mers on the distance distribution between selected positions we used 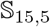(3,9) on the unmutated CHM13X. Fig S2A shows the distance distribution for syncmers selected only using forward strand *k*-mers. The minimum distance is 3 and there is a sharp peak at 6. In general, for a scheme with parameters *x*_1_ <… < *x_n_*, the minimum distance is *min*(*x*_1_, *k* – *s* – *x_n_* + 2) and there are peaks at *x*_*i*+1_ – *x_i_*, in agreement with the expression for *D*(*α*) above (Calculating *ℓ*_2,*mut*_). In mapping, read orientations are unknown and canonical syncmers are used. Fig S2B shows the results using canonical *k*-mers.

**Table S2.** Properties of minimizer and syncmer schemes on real sequences without mutations. The best syncmer schemes with theoretical compression of 5.5 and 10 were chosen. The table shows the actual compression and other metrics on the real sequences, # positions is the number of positions that were selected by the scheme.

| Dataset | Scheme | Compression | $\ell$ | $\ell_2$ | $p_{90}$ | $p_{100}$ | # positions |
| --- | --- | --- | --- | --- | --- | --- | --- |
| ECK12 | $\mathcal{M}_{15,10}$ | 5.503 | $3.23 \times 10^{-6}$ | 0.005 | 9 | 10 | 843,408 |
| | $\mathbb{S}_{15,5}(3, 9)$ | 5.490 | 0.0191 | 0.377 | 10 | 53 | 845,419 |
| | $\mathbb{S}_{15,5}(1, 11)$ | 5.509 | 0.0255 | 0.443 | 11 | 52 | 842,557 |
| | $\mathbb{S}_{15,5}(1, 2)$ | 5.502 | 0.0566 | 0.715 | 13 | 66 | 843,634 |
| | $\mathcal{M}_{15,19}$ | 9.989 | 0.0527 | 0.398 | 18 | 19 | 464,660 |
| | $\mathbb{S}_{15,6}(6)$ | 9.986 | 0.133 | 1.375 | 19 | 99 | 464,810 |
| CHM13X | $\mathcal{M}_{15,10}$ | 5.489 | $5.83 \times 10^{-8}$ | 0.0005 | 9 | 10 | 28,099,399 |
| | $\mathbb{S}_{15,5}(3, 9)$ | 5.523 | 0.0205 | 0.412 | 10 | 161 | 27,930,897 |
| | $\mathcal{M}_{15,19}$ | 9.977 | 0.0526 | 0.3974 | 18 | 19 | 15,461,458 |
| | $\mathbb{S}_{15,6}(6)$ | 10.055 | 0.137 | 1.419 | 20 | 735 | 15,342,238 |
| | $\mathbb{S}_{15,5}(1, 11)$ | 5.437 | 0.0259 | 0.449 | 11 | 260 | 28,371,548 |
| | $\mathbb{S}_{15,5}(1, 2)$ | 5.350 | 0.0549 | 0.706 | 13 | 122 | 28,835,008 |

The distance distribution still retains the peak at 6 and a local maximum at 3, but now adjacent positions are selected, and it has a much longer tail of distances (compare Supplementary Table S2). We conclude that while the theory is limited to single-stranded sequences it shows trends that hold for canonical *k*-mers.

### S9 Supplemental performance results

Fig S3 shows additional results for the number of unmapped reads at low, medium, and high compression rates.

Fig S4 compares the performance of the theoretically optimal 2-parameter PSS and closed syncmers in mapping real PacBio bacterial reads against the BAC reference.

### S10 Windowed syncmer scheme results

Tables S3 and S4 present the properties of windowed syncmer schemes on real genome sequences with and without mutation, respectively.

Figures S5 and S6 present the number of unmapped reads and wrongly mapped reads for simulated datasets. These correspond to Figure 4 and include the results for windowed variants. Fig S7 presents the impact of percent sequence identity on the windowed variants as well, corresponding to Fig 6.

Results on the real human and bacterial reads are presented in Fig S8, and the runtimes and RAM usage for these runs are in Figures S9 and S10. The runtime and memory usage on different tasks for the windowed variants is presented in Table S5.

**Fig S2.**
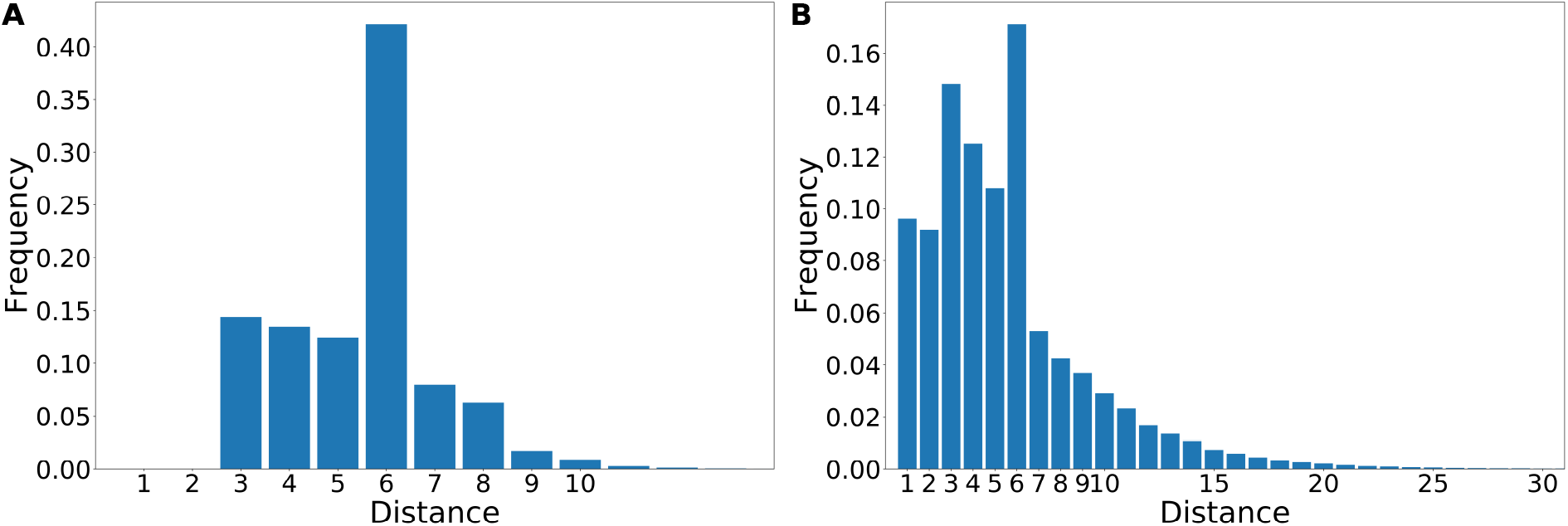
Distribution of the distances between selected positions in a syncmer scheme. The distribution of distances between consecutive selected positions of syncmer scheme 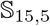(3, 9) on the CHM13X reference is shown. **A.** The distribution of syncmers selected only in the forward orientation. **B.** Canonical syncmers. For visualization purposes the distribution is shown only for distances with frequency > 10^-5^. The true maximum distance is 161 for canonical *k*-mers (see Supplementary Table S2) and 76 for the forward *k*-mers, but the frequency of the longer distances is extremely low.

**Fig S3.**
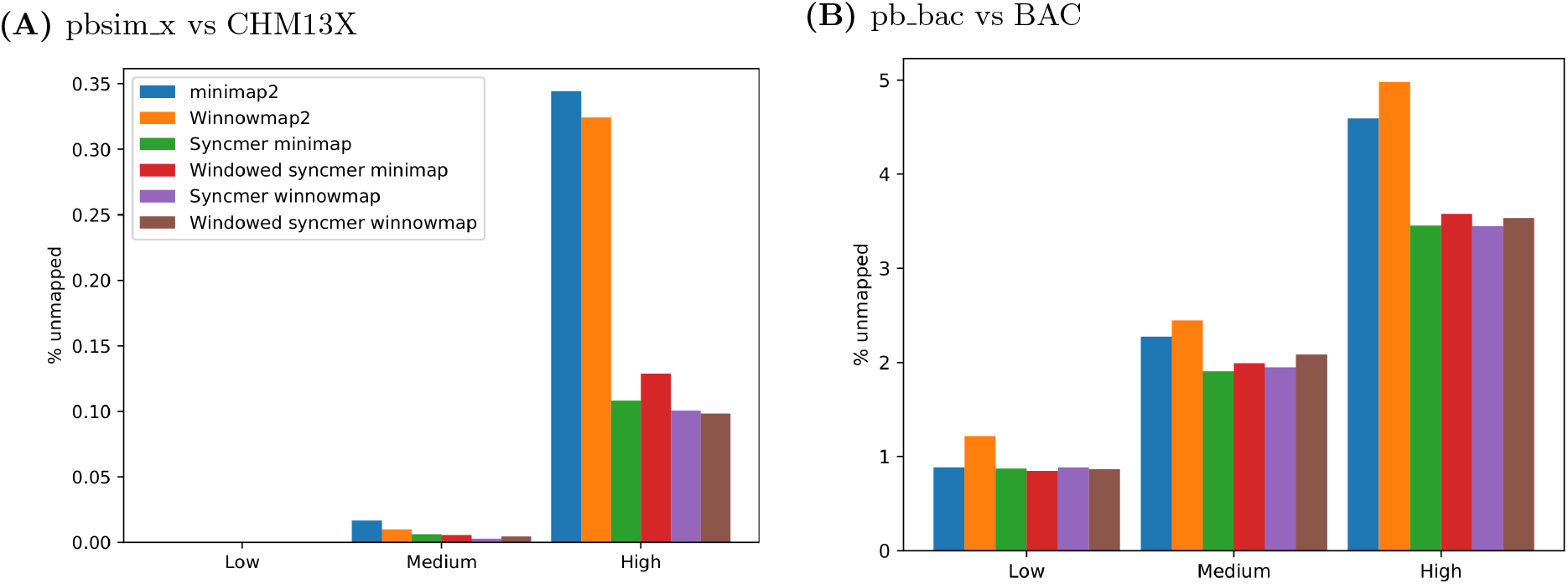
Percentage of unmapped reads – additional data. The percentage of unmapped reads is plotted for one simulated and one real read dataset mapped to their corresponding references. Minimizer and PSS parameters are as described in the Results section (The fraction of unmapped reads). **(A)** PacBio reads simulated from the CHM13 ChrX sequence mapped against CHM13X. Window sizes of windowed syncmer-minimap were *w* = 13, 80, 165 for the low, medium, and high compression variants, respectively, and for windowed syncmer-winnowmap they were *w* = 14, 75, 170, respectively. **(B)** 1000 PacBio reads sampled from each of the 15 bacterial species in BAC mapped against their reference genomes. Window sizes of windowed syncmer-minimap were *w* = 13, 75, 175 and in windowed syncmer-winnowmap they were *w* = 16, 75, 170 for the low, medium, and high compression variants, respectively.

**Fig S4.**
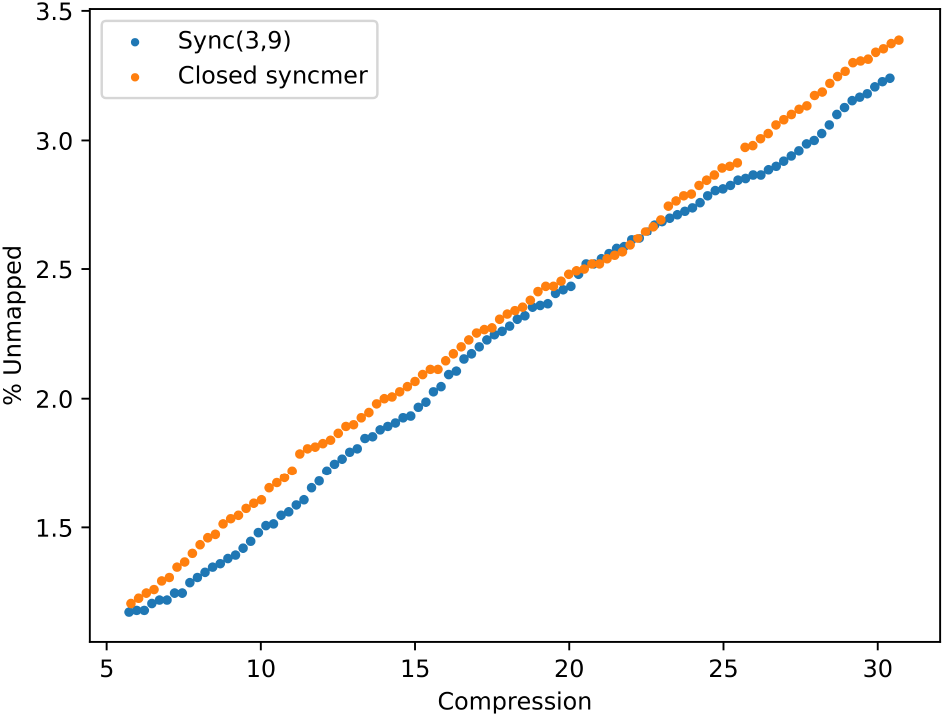
Comparison of 2-parameter PSS to closed syncmers. 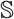(3, 9) and closed syncmers were used in syncmer-minimap to map PacBio bacterial reads against BAC. Both schemes used *k* = 15, *s* = 5.

**Table S3.**
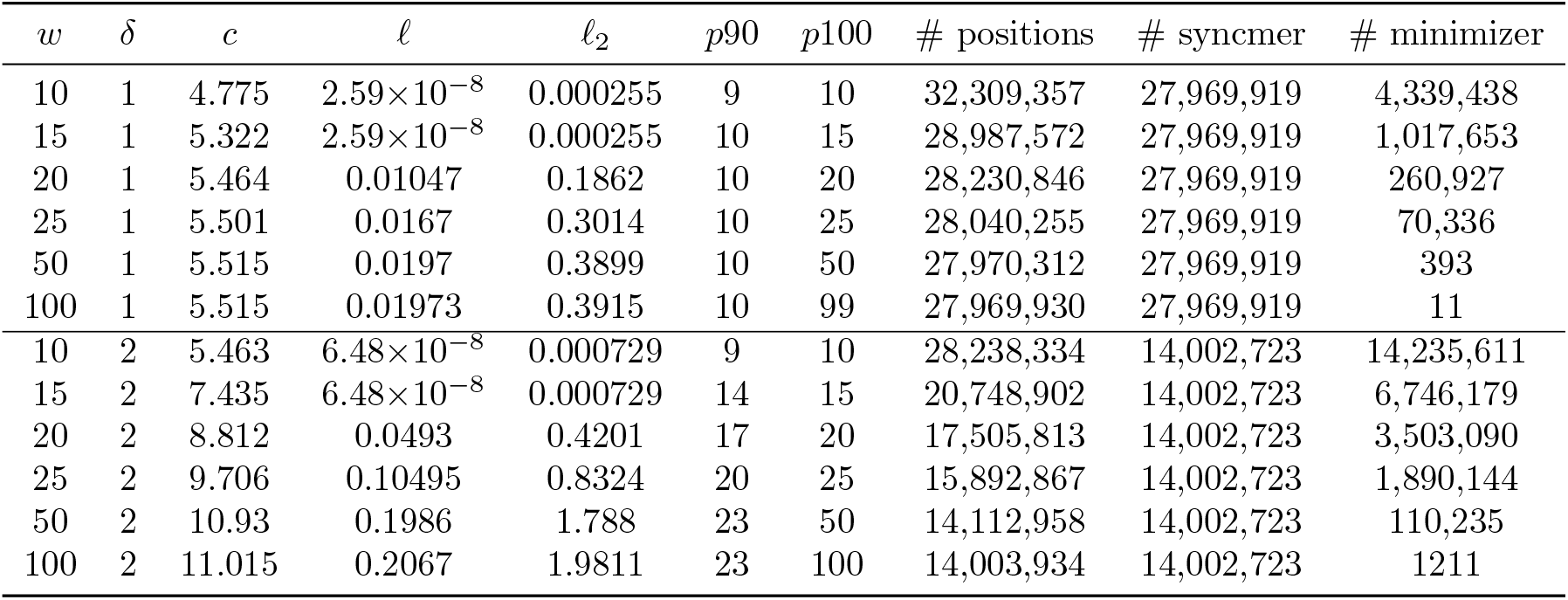
Properties of windowed syncmers on CHM13X. Properties of windowed and down-sampled variants of 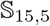(3, 9) are shown for a range of window lengths w on the human chromosome X sequence of CHM13. *δ* is the downsampling rate and *c* is the actual compression. The theoretical compression of the PSS (not downsampled) is 5.5. # positions is the number of positions that were selected by the scheme, # syncmers is the number of those that were selected by the PSS and # minimizers is the number of minimzers added to fill in gaps of length ≥ *w*.

**Table S4.** Properties of conserved windowed syncmers under mutation. Properties of windowed and down-sampled variants of 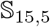(3, 9) are shown for a range of window lengths *w* on CHM13X after simulating substitutions at a rate of 15%. w is the window size and δ is the downsampling rate. Properties of the conserved selected *k*-mers are reported.

| $w$ | $\delta$ | $c$ | $\ell$ | $\ell_2$ | $p90$ | $p100$ | # conserved | conserved syncmers | conserved minimizers |
| --- | --- | --- | --- | --- | --- | --- | --- | --- | --- |
| 10 | 1 | 41.745 | 0.7826 | 10.713 | 122 | 1221 | 3,695,319 | 3,378,302 | 317,017 |
| 15 | 1 | 45.008 | 0.7928 | 11.041 | 131 | 1221 | 3,427,368 | 3,378,302 | 49066 |
| 20 | 1 | 45.558 | 0.7951 | 11.1125 | 132 | 1221 | 3,386,019 | 3,378,302 | 7717 |
| 25 | 1 | 45.646 | 0.7955 | 11.126 | 132 | 1221 | 3,379,466 | 3,378,302 | 1164 |
| 50 | 1 | 45.662 | 0.7956 | 11.129 | 133 | 1274 | 3,378,304 | 3,378,302 | 2 |
| 100 | 1 | 45.662 | 0.7956 | 11.128 | 133 | 1274 | 3,378,302 | 3,378,302 | 0 |
| 10 | 2 | 52.34 | 0.8062 | 11.515 | 147 | 1221 | 2,947,249 | 1,692,160 | 1,255,089 |
| 15 | 2 | 71.027 | 0.8397 | 12.898 | 190 | 1691 | 2,171,847 | 1,692,160 | 479,687 |
| 20 | 2 | 81.734 | 0.8572 | 13.774 | 216 | 1703 | 1,887,340 | 1,692,160 | 195,180 |
| 25 | 2 | 86.999 | 0.8655 | 14.257 | 230 | 1736 | 1,773,121 | 1,692,160 | 80961 |
| 50 | 2 | 91.104 | 0.8722 | 14.709 | 242 | 1736 | 1,693,222 | 1,692,160 | 1062 |
| 100 | 2 | 91.161 | 0.8723 | 14.718 | 242 | 1736 | 1,692,160 | 1,692,160 | 0 |

**Fig S5.**
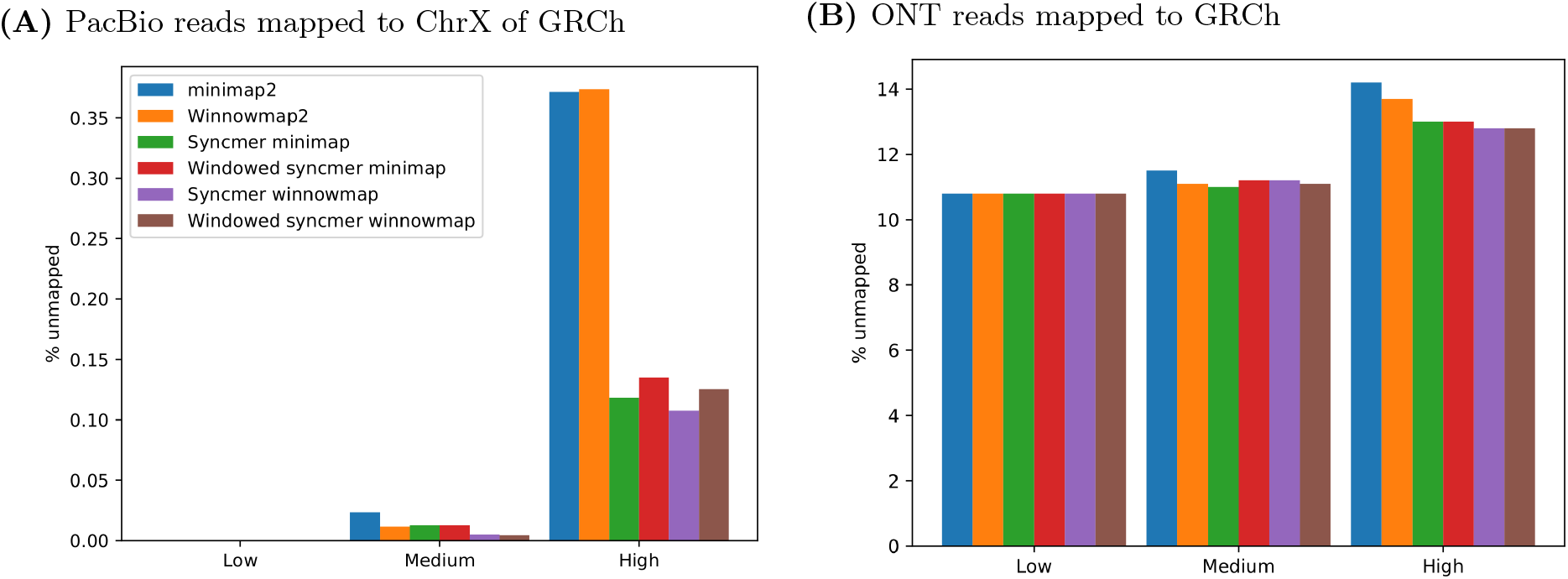
Percentage of unmapped reads – simulated datasets. The percentage of unmapped reads is plotted for two simulated read datasets mapped to their reference sequences. Results are shown for low, medium, and high compression. **(A)** PacBio reads simulated from the CHM13 ChrX sequence mapped against ChrX sequences from GRCh38. Window sizes of windowed syncmer-minimap were *w* = 13, 77, 175 for the low, medium, and high compression variants, respectively. For windowed syncmer-winnowmap the window sizes were *w* = 14, 75, 170, respectively. **(B)** 1000 ONT reads simulated from CHM13 mapped against GRCh38. Window sizes of windowed syncmer-minimap were *w* = 13, 75, 175 for the low, medium, and high compression variants, respectively and *w* = 13, 75, 170 for the corresponding windowed syncmer-winnowmap variants.

**Fig S6.**
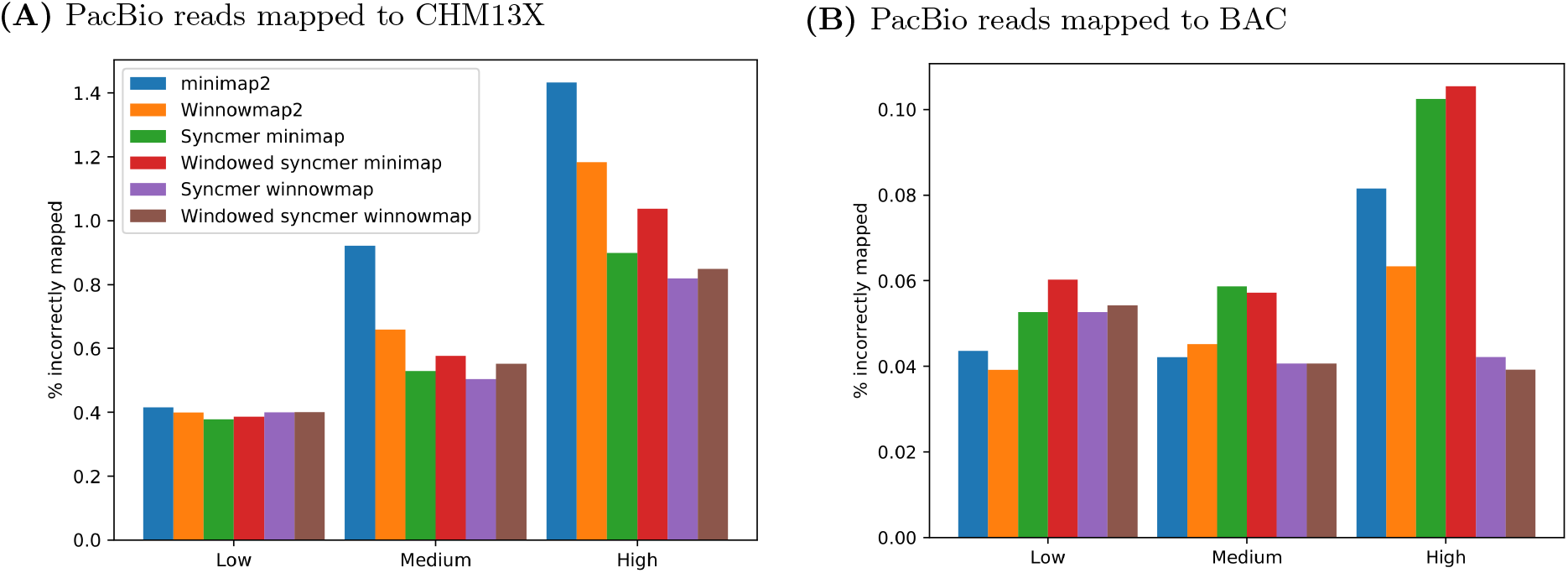
Percentage of incorrectly mapped reads – simulated data. The percentage of incorrectly mapped reads is plotted for two simulated read datasets and their reference sequences, for mappers using low, medium, and high compression. **(A)** PacBio reads simulated from the CHM13 ChrX sequence mapped against CHM13X. Window sizes of windowed syncmer-minimap were *w* = 13, 80, 165 for the low, medium, and high compression variants, respectively. For windowed syncmer-winnowmap they were *w* = 14, 75, 170, respectively. **(B)** PacBio reads simulated from the 15 bacterial species in BAC mapped against the union of their references. Window sizes of windowed syncmer-minimap were *w* = 13, 75, 175 and in windowed syncmer-winnowmap they were *w* = 16, 75, 170 for the low, medium, and high compression variants, respectively.

**Fig S7.**
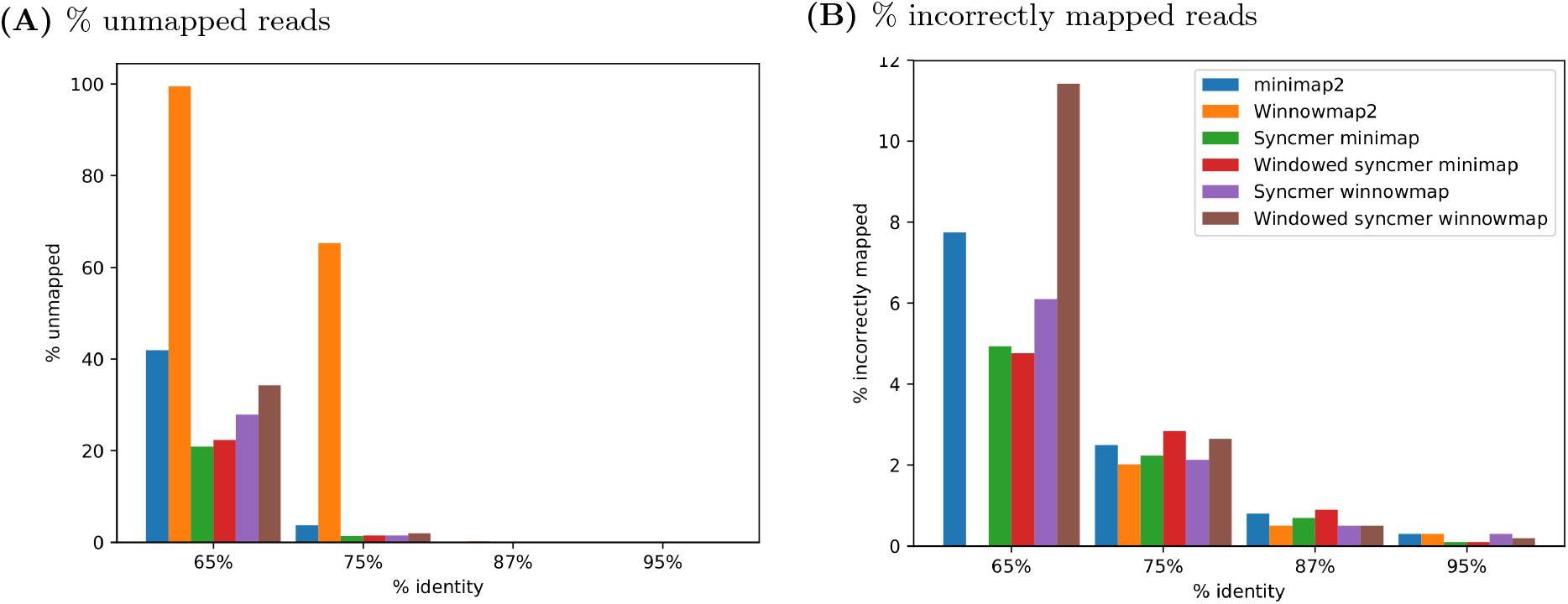
Impact of percent sequence identity. We varied the mutation rate of 1000 PacBio simulated reads from CHM13X. The figures present the % unmapped and incorrectly mapped for each of the tools. **(A)** % unmapped reads. **(B)** % of the mapped reads that were incorrectly mapped.

**Fig S8.**
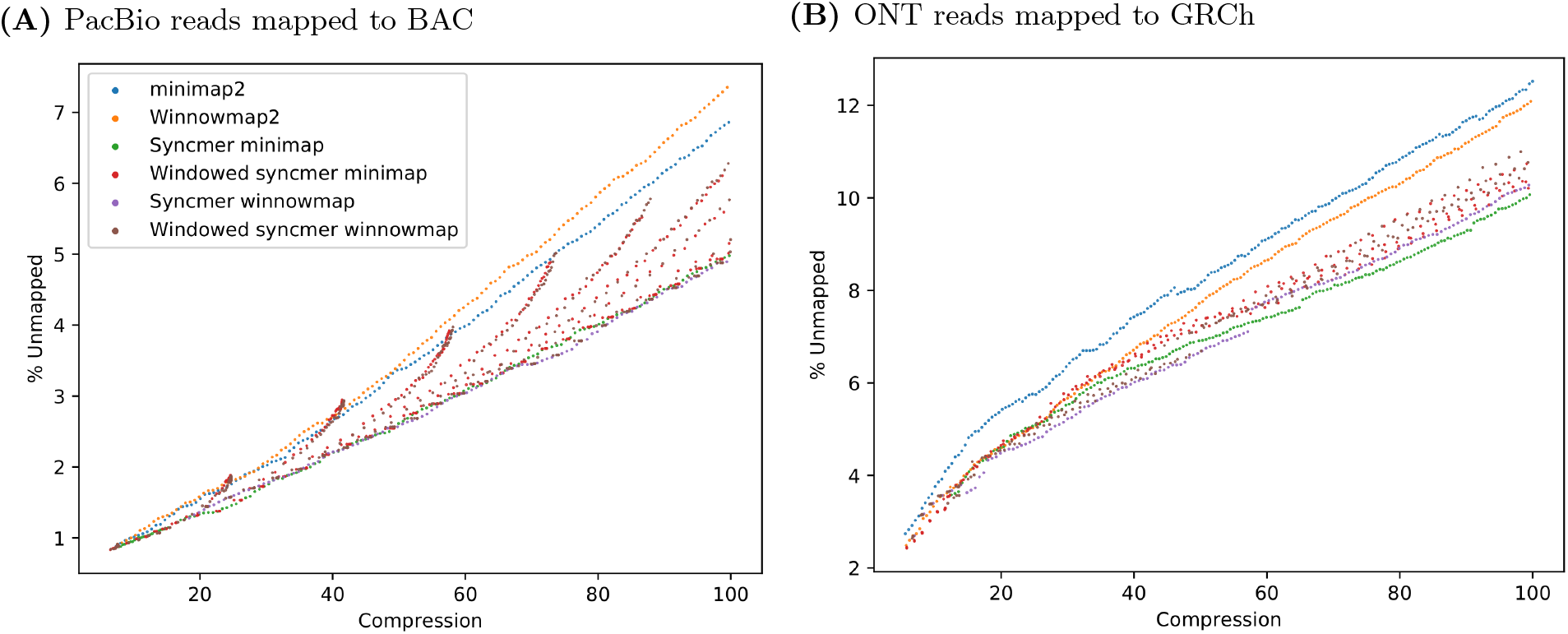
Percentage of unmapped reads – real datasets. Results are shown across a broad range of compression rates. **(A)** Pooled PacBio bacterial reads. **(B)** ONT human cell-line reads.

**Fig S9.**
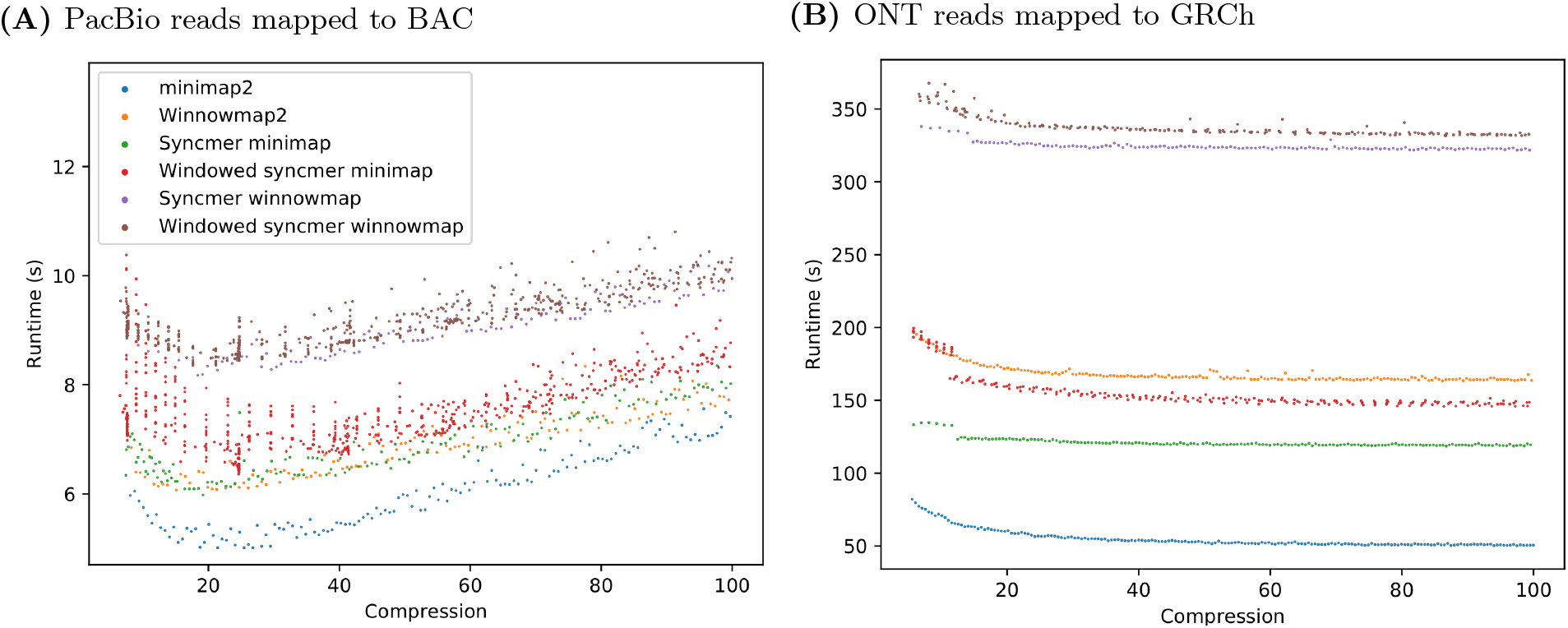
Runtime vs. compression – real data. The figures show runtime in seconds to index the reference and map reads by each method. **(A)** PacBio bacterial reads. **(B)** ONT human cell-line reads.

**Fig S10.**
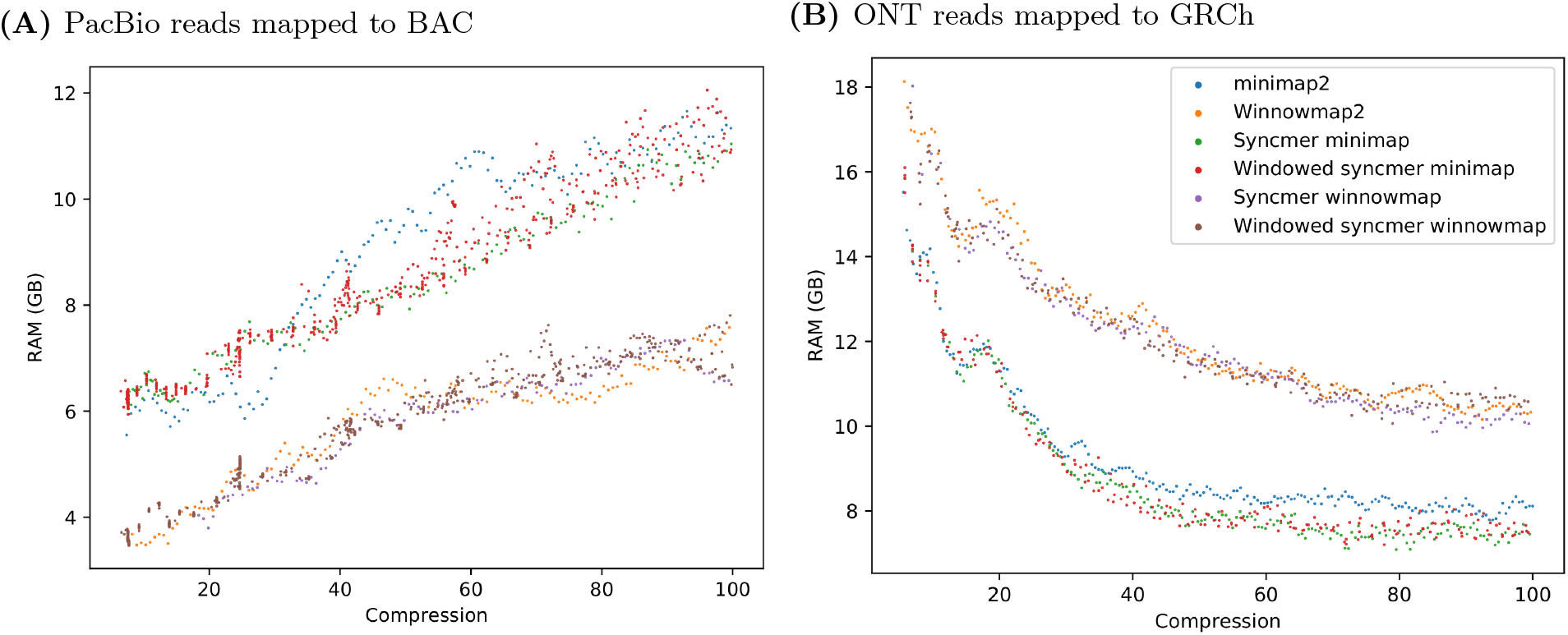
Memory usage vs. compression – real data. Peak RAM usage in GB to index the reference and map reads for the different methods. **(A)** PacBio bacterial reads. **(B)** ONT human cell-line reads.

**Table S5.**
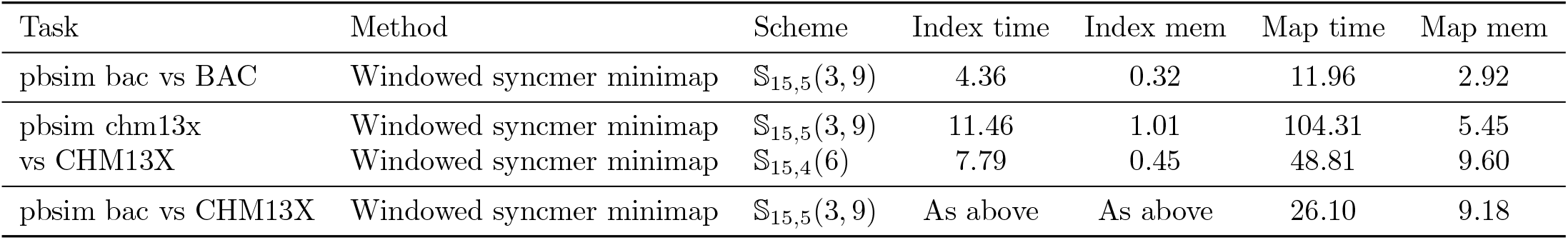
Runtime and memory. Time (in seconds) and RAM (in GB) needed to index the reference and map reads by each of the tools. The second and third dataset use the same reference and therefore have the same indexing results. For 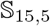(3, 9) window size *w* = 13 was used, with downsampling rate of 1.08 for BAC and downsampling rate of 1.05 for CHM13X. For 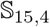(6) the standard PSS used downsampling rate of 4.13, and the windowed PSS had window size *w* = 165 and downsampling rate 4.37.

## References

1. Jain C, Rhie A, Zhang H, Chu C, Walenz BP, Koren S, et al. Weighted minimizer sampling improves long read mapping. Bioinformatics. 2020;36(Supplement_1):i111–i118.

2. Sedlazeck FJ, Rescheneder P, Smolka M, Fang H, Nattestad M, Von Haeseler A, et al. Accurate detection of complex structural variations using single-molecule sequencing. Nature methods. 2018;15(6):461–468.

3. Li H. Minimap2: pairwise alignment for nucleotide sequences. Bioinformatics. 2018;34(18):3094–3100.

4. Chikhi R, Limasset A, Medvedev P. Compacting de Bruijn graphs from sequencing data quickly and in low memory. Bioinformatics. 2016;32(12):i201–i208.

5. Kokot M, Długosz M, Deorowicz S. KMC 3: counting and manipulating k-mer statistics. Bioinformatics. 2017;33(17):2759–2761.

6. Wood DE, Lu J, Langmead B. Improved metagenomic analysis with Kraken 2. Genome biology. 2019;20(1):1–13.

7. Edgar R. Syncmers are more sensitive than minimizers for selecting conserved k-mers in biological sequences. PeerJ. 2021;9:e10805.

8. Dohm JC, Peters P, Stralis-Pavese N, Himmelbauer H. Benchmarking of long-read correction methods. NAR Genomics and Bioinformatics. 2020;2(2):lqaa037.

9. Shaw J, Yu YW. Theory of local k-mer selection with applications to long-read alignment. Bioinformatics. 2021;doi:10.1093/bioinformatics/btab790.

10. Li H. New strategies to improve minimap2 alignment accuracy. arXiv preprint arXiv:210803515. 2021;.

11. Jain C, Rhie A, Hansen NF, Koren S, Phillippy AM. Long-read mapping to repetitive reference sequences using Winnowmap2. Nature Methods. 2022; p. 1–6.

12. Schleimer S, Wilkerson DS, Aiken A. Winnowing: local algorithms for document fingerprinting. In: Proceedings of the 2003 ACM SIGMOD international conference on Management of data; 2003. p. 76–85.

13. Schneider VA, Graves-Lindsay T, Howe K, Bouk N, Chen HC, Kitts PA, et al. Evaluation of GRCh38 and de novo haploid genome assemblies demonstrates the enduring quality of the reference assembly. Genome research. 2017;27(5):849–864.

14. Nurk S, Koren S, Rhie A, Rautiainen M, Bzikadze AV, Mikheenko A, et al. The complete sequence of a human genome. Science. 2022;376(6588):44–53. doi:10.1126/science.abj6987.

15. Blattner FR, Plunkett G, Bloch CA, Perna NT, Burland V, Riley M, et al. The complete genome sequence of Escherichia coli K-12. Science. 1997;277(5331):1453–1462.

16. PacificBiosciences. Microbial Multiplexing Data Set 48 plex: PacBio Sequel II System, Chemistry v2.0, SMRT Link v8.0 Analysis; 2019. https://github.com/PacificBiosciences/DevNet/wiki/Microbial-Multiplexing-Data-Set---48-plex:-PacBio-Sequel-II-System,-Chemistry-v2.0,-SMRT-Link-v8.0-Analysis.

17. Ono Y, Asai K, Hamada M. PBSIM: PacBio reads simulator—toward accurate genome assembly. Bioinformatics. 2013;29(1):119–121.

18. Yang C, Chu J, Warren RL, Birol I. NanoSim: nanopore sequence read simulator based on statistical characterization. GigaScience. 2017;6(4):gix010.

## References

1. Shaw J, Yu YW. Theory of local k-mer selection with applications to long-read alignment. Bioinformatics. 2021;doi:10.1093/bioinformatics/btab790.

2. Ono Y, Asai K, Hamada M. PBSIM: PacBio reads simulator—toward accurate genome assembly. Bioinformatics. 2013;29(1):119–121.

3. Dohm JC, Peters P, Stralis-Pavese N, Himmelbauer H. Benchmarking of long-read correction methods. NAR Genomics and Bioinformatics. 2020;2(2):lqaa037.

4. Yang C, Chu J, Warren RL, Birol I. NanoSim: nanopore sequence read simulator based on statistical characterization. GigaScience. 2017;6(4):gix010.

5. Jain C, Rhie A, Zhang H, Chu C, Walenz BP, Koren S, et al. Weighted minimizer sampling improves long read mapping. Bioinformatics. 2020;36(Supplement_1):i111–i118.

6. PacificBiosciences. Microbial Multiplexing Data Set 48 plex: PacBio Sequel II System, Chemistry v2.0, SMRT Link v8.0 Analysis; 2019. https://github.com/PacificBiosciences/DevNet/wiki/Microbial-Multiplexing-Data-Set---48-plex:-PacBio-Sequel-II-System,-Chemistry-v2.0,-SMRT-Link-v8.0-Analysis.

